# Preclinical Evaluation of a Novel BOC-Modified Triazoloacridinone (TRA-01AR) in Combination with 5-Fluorouracil in Colorectal Cancer

**DOI:** 10.64898/2026.11.08.765151

**Authors:** Marta Świtalska, Paulina Czechowicz, Magdalena Milczarek, Sylwia Bartoszewska, Beata Filip-Psurska, Katarzyna Wierzbicka, Mateusz Psurski, Damian Szczesny, Eliza Turlej, Karol Krzymiński, Małgorzata Gawrońska, Magdalena Gebert, Leszek Kalinowski, David K. Crossman, Maciej Bagiński, Michał Markuszewski, James F. Collawn, Liliana Schaefer, Rafal Bartoszewski, Joanna Wietrzyk

## Abstract

5-Fluorouracil (5-FU) remains a cornerstone of colorectal cancer (CRC) therapy, yet its clinical efficacy is limited by resistance, systemic toxicity, and unfavorable pharmacokinetics. Strategies that enhance tumor sensitivity while maintaining tolerability are therefore of considerable interest. Here, we evaluated 5-FU in combination with two triazoloacridinones, C-1305 and a novel BOC-modified derivative TRA-01AR, using integrated cellular, transcriptomic, bioenergetic, and *in vivo* approaches. Both compounds induced cytotoxic effects across cancer cell models and produced overlapping but distinct transcriptional responses, primarily affecting DNA replication and the cell cycle pathways. Bioenergetic profiling revealed a shift toward oxidative phosphorylation accompanied by reduced glycolysis, indicating modulation of cellular metabolism relevant to the chemotherapy response. Interestingly, TRA-01AR exhibited a delayed cytotoxicity relative to C-1305, although the underlying mechanism remains to be defined. *In vivo*, the combination of 5-FU with C-1305 significantly reduced tumor weight in HT-29 xenografts, whereas 5-FU combined with TRA-01AR did not result in a statistically significant reduction compared to 5-FU alone. More importantly, TRA-01AR was associated with a more favorable systemic profile and increased intratumoral blood flow compared to C-1305. Collectively, these findings indicate that triazoloacridinones modulate transcriptional and metabolic programs relevant to the 5-FU response. While C-1305 demonstrates a higher antitumor activity under the tested conditions, TRA-01AR exhibits distinct biological effects, including improved tolerability and vascular modulation. Further studies are required to clarify TRA-01AR’s mechanism of action and assess its potential relevance in CRC-specific therapies.

**Mesh Terms:** Fluorouracil; Colorectal Neoplasms/drug therapy; Drug Resistance, Neoplasm; Drug Synergism; Acridines/pharmacology; Prodrugs/pharmacology; Oxidative Phosphorylation; Tumor Microenvironment; Angiogenesis Inhibitors/pharmacology; Xenograft Model Antitumor Assays; Transcriptome; Gene Expression Profiling; RNA Sequencing; Gene Expression Regulation, Neoplastic; High-Throughput Nucleotide Sequencing; Bioinformatics; Signal Transduction; Apoptosis/drug effects; Mitochondria/metabolism; Cell Cycle/drug effects

## INTRODUCTION

5-Fluorouracil (5-FU) remains a central component of colorectal cancer (CRC) therapy, forming the backbone of widely used regimens such as FOLFOX, FOLFIRI and FOLFOXIRI [1]. Its cytotoxic activity derives primarily from thymidylate synthase inhibition and incorporation of fluorinated nucleotides into RNA and DNA and leads to impaired nucleic acid synthesis and replication stress [2–5]. Despite its long-standing clinical utility, the therapeutic benefit of 5-FU is frequently limited by the emergence of resistance, systemic toxicity, and unfavorable pharmacokinetic properties that include rapid catabolism and a short plasma half-life [2, 3, 6]. Contemporary strategies to overcome these limitations increasingly rely on combination approaches that target complementary vulnerabilities such as metabolic dependencies, DNA damage responses, and tumor microenvironment dynamics [7].

Resistance to 5-FU has been associated with metabolic reprogramming, particularly a shift toward increased oxidative phosphorylation (OXPHOS), which supports survival under chemotherapeutic stress [8, 9]. Such bioenergetic adaptations provide a rationale for combining 5-FU with agents that perturb mitochondrial function or redox balance [10–12]. In parallel, modulation of tumor perfusion and vascular function has emerged as an additional determinant of therapeutic efficacy [13, 14] by influencing drug delivery and intratumoral distribution [15, 16]. Together, these insights highlight the need for combination partners that not only enhance cytotoxicity but also improve tumor accessibility and systemic tolerability.

Triazoloacridinones constitute a class of antitumor compounds with pleiotropic mechanisms of action, including DNA intercalation, topoisomerase II inhibition, and microtubule stabilization [17–25]. Among these, C-1305 (**Figure 1A**) has demonstrated potent antitumor activity in preclinical models, including CRC [26, 27]. Its development, however, has been limited by its toxicity and the incomplete characterization of its structure–activity relationships [22, 28]. Related compounds such as C-1311 (Symadex™, NSC-645809) have progressed to early clinical evaluation, which supports the translational potential of this chemical scaffold [29, 30] (**Figure 1B**). Previous studies have also suggested that triazoloacridinones influence tumor angiogenesis and perfusion when used in combination regimens, which also indicates their relevance as partners in multi-modal anticancer strategies [28, 29, 31].

**Figure 1.**
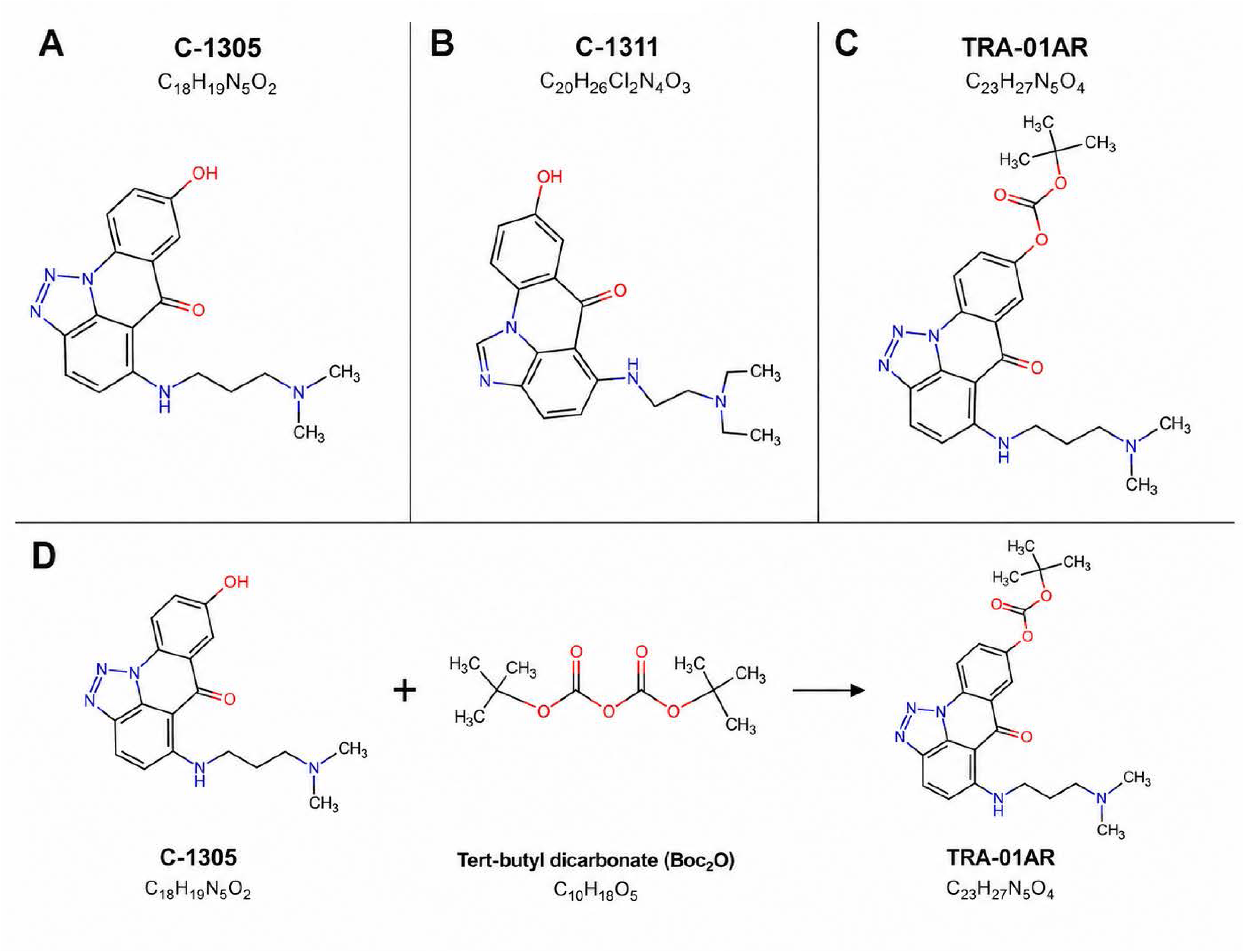
Chemical structures of triazoloacridinones and synthesis of TRA-I01AR. (**A**) C-1305, (**B**) C-1311, and (**C**) TRA-01AR. (**D**) Reaction scheme for the synthesis of TRA-01AR via BOC modification of C-1305.

Based on these studies, we developed TRA-01AR, a novel tert-butyloxycarbonyl (BOC)-modified derivative of C-1305 (**Figure 1C**). The introduction of the BOC group was designed to alter key physicochemical properties, including lipophilicity and ionization state, with the goal of modulating the cellular uptake and intracellular engagement. Such modifications are conceptually consistent with the prodrug-oriented design strategies that are intended to improve physicochemical and pharmacokinetic properties [32, 33]. Direct evidence of intracellular deprotection and activation, however, remains to be established. Notably, preliminary observations indicate that TRA-01AR exhibits a delayed cytotoxic effect compared to C-1305, suggesting that the differences in cellular processing may influence both efficacy and tolerability.

Our previous studies using triazoloacridinones demonstrated that C-1305 exerts pleiotropic effects on cellular processes that include direct microtubule stabilization identified by genome-wide mRNA profiling, modulation of the IRE1α/XBP1 signaling pathway, and tumor angiogenesis in *in vivo* models [22, 25, 28]. These findings established a foundation for further mechanistic investigation using high-resolution transcriptomic approaches.

We hypothesized that triazoloacridinones through their induction of bioenergetic and transcriptional reprogramming could enhance the therapeutic response to 5-FU in CRC. To test this, we evaluated the combination of 5-FU with C-1305 and with TRA-01AR using a multi-level approach that integrated cytotoxicity assays in CRC models, transcriptomic profiling, bioenergetic analysis, and *in vivo* validation in HT-29 xenografts. Mechanistic studies were initially performed in a well-characterized epithelial cancer model to enable high-resolution analysis of drug-induced transcriptional responses that were validated in CRC-relevant systems.

Here, we demonstrate that both triazoloacridinones induce distinct but partially overlapping transcriptional and metabolic responses that include modulation of DNA replication pathways and bioenergetic shifts toward oxidative phosphorylation. While combination therapy with 5-FU and C-1305 significantly reduced tumor burden *in vivo*, TRA-01AR exhibited a non-significant trend toward tumor reduction but importantly, demonstrated a more favorable systemic profile and enhanced tumor perfusion. These findings justify further investigation of TRA-01AR to clarify its mechanism of action, optimize dosing strategies, and assess its potential relevance in combination therapies with 5-FU.

## RESULTS

### Synthesis and characterization of TRA**-I**01AR

The synthesis of TRA-01AR was carried out using purified C-1305 as the starting material (see Methods section). C-1305 was dissolved in dichloromethane (CH₂Cl₂), and a five-fold molar excess of triethylamine and di-tert-butyl dicarbonate was added. The reaction mixture was stirred at room temperature for 2 h. Upon completion, the solvent and residual triethylamine were removed under reduced pressure. The resulting residue was purified by flash chromatography on neutral alumina (Al₂O₃) using a CHCl₃/hexane (1:1, v/v) eluent system since preliminary studies demonstrated instability of the compound on silica gel. The purified product was obtained as a solid powder with a synthetic yield of 69%. The reaction scheme is shown in **Figure 1D**.

The obtained compound was characterized by high-resolution mass spectrometry (HRMS), RP-HPLC, melting point analysis, and proton nuclear magnetic resonance (^1H NMR) spectroscopy. HRMS analysis revealed a molecular ion peak at m/z 438.2148, consistent with the calculated value of 438.2261. RP-HPLC analysis indicated a chromatographic purity of 94.52%. The ^1H NMR spectrum displayed characteristic signals corresponding to the aromatic acridone core, methylene linker, and *tert*Rlbutyl moiety, with signal assignments supported by COSY correlations. Collectively, HRMS, RP-HPLC, melting point analysis, and NMR spectroscopy support the structural assignment and analytical characterization of TRA-01AR (**Supplementary Figure 1)**.

### Cytotoxic effects of C**-I**1305 and TRA**-I**01AR in cancer and non**-**cancer cell lines

To compare the cytotoxicity of C-1305 and TRA-01AR, MTT assays were performed on cancer and normal human cell lines previously used in studies of C-1305 activity [34, 35]. In A549 non-small cell lung cancer cells, TRA-01AR exhibited a significantly higher IC₅₀ at 24 h (11.78 µM, p = 0.001) compared to C-1305 (3.08 µM, p = 0.0024). However, at 48 h TRA-01AR demonstrated a lower IC₅₀ value (IC₅₀ = 0.942 µM, p = 0.002) than C-1305 (IC₅₀ = 1.900 µM, p = 0.002), indicating increased cytotoxic potency. In HCT-116 CRC cells, 24 h IC₅₀ values were 9.27 µM for C-1305 and 18.68 µM for TRA-01AR, while at 48 h they were 9.01 µM and 12.13 µM, respectively.

In the immortalized normal human bronchial epithelial cell line (16HBE14o-), C-1305 showed IC₅₀ values of 23.66 µM (24 h) and 31.19 µM (48 h), whereas TRA-01AR exhibited no toxicity up to 50 µM at 24 h and an IC₅₀ of 31.12 µM at 48 h. Notably, concentrations below 5 µM of either compound had no significant effect on 16HBE14o-cell growth.

Additionally, the antiproliferative activity of C-1305 and TRA-01AR (72 h, SRB assay) was compared with 5-FU in colon cancer cell lines, fetal human colonic (FHC) cells, and human lung microvascular endothelial cells (HLMEC). Both triazoloacridinones were more effective than 5-FU against selected colon adenocarcinoma cell lines (HCT-116, SW480, LS180, HT-29) but also showed high cytotoxicity toward normal FHC and HLMEC cells. All three agents displayed strong activity (IC₅₀ < 5 μM) against additional colon cancer cell lines (LoVo, SW620, LS1034, DLD1) (**Supplementary Figure 2**).

All cytotoxicity data are summarized in **Table 1** and **Supplementary Figure 2**. These results guided dose selection for subsequent RNA-seq experiments and *in vivo* studies.

**Table 1.** Cytotoxicity of tested compounds towards normal and cancer cell lines expressed as IC₅₀ values (µM). Data are presented as mean ± SD.

| **Cell Line** | **Compound/IC<sub>50</sub> [µM] mean ± SD** |  |  |
|---|---|---|---|
|  | **C-1305** | **TRA01AR** | **5-FU** |
| **FHC** | 3.6 ± 0.4* | 1.6 ± 0.4* | 11.2 ± 9.2 |
| **16HBE14o** | 31.19 ± 0.5 | 31.12 ± 1.1 | ND |
| **HLMEC** | 5.8 ± 0.8* | 6.0 ± 0.4* | 31.9 ± 11.2 |
| **DLD-1** | 1.3 ± 0.7 | 1.2 ± 0.5 | 0.8 ± 0.8 |
| **LoVo** | 1.6 ± 0.1 | 1.3 ± 0.4 | 3.1 ± 0.2 |
| **HT-29** | 6.9 ± 1.9 | 5.8 ± 0.4 | 12.6 ± 4.1 |
| **HCT-116** | 6.7 ± 1.5 | 6.1 ± 1.2* | 13.5 ± 8.5 |
| **LS180** | 1.9 ± 0.3* | 2.0 ± 0.7* | 24.0 ± 3.5 |
| **LS1034** | 4.3 ± 0.1 | 3.7 ± 0.7 | 0.6 ± 0.1 |
| **SW-480** | 5.5 ± 1.1* | 5.6 ± 1.0* | 39.1 ± 9.7 |
| **SW-620** | 1.1 ± 0.3 | 1.5 ± 0.2 | 0.6 ± 0.3 |
| **A549** | 1.9 ± 0.14 | 0.94 ± 0.1 | ND |

N=3-4 with three technical replicates in each biological replicate. Statistical analysis: two-way ANOVA revealed a significant interaction between cell line and compound (F(18,59) = 12.83, p < 0.0001), indicating that the effects of the compounds differed among the cell lines. Significant main effects of cell line (F(9,59) = 22.25, p < 0.0001) and compound (F(2,59) = 81.37, p < 0.0001) were also observed. Šídák’s multiple comparisons test: \*p < 0.05, \*\*p < 0.01, \*\*\*p < 0.001, \*\*\*\*p < 0.0001 compared with 5-FU for each cell line. ND - not determined

### Selection of experimental conditions for transcriptomic analysis

Based on cytotoxicity data, A549 cells were selected for transcriptomic analysis due to consistent responses across treatment conditions. Cells were treated with C-1305 or TRA-01AR at concentrations corresponding to their IC₅₀ values (3 µM C-1305 or 2 µM TRA-01AR) and harvested after 24 h and 48 h. Vehicle (DMSO)-treated controls were collected at matching time points (24 h and 48 h). Non-cancerous 16HBE14o⁻ cells were included as a reference for conditions in which limited cytotoxicity was observed. To account for the potential genomic instability reported in A549 cells [16] and to control for time-dependent transcriptional changes unrelated to compound exposure, all transcriptomic analyses were performed using time-matched vehicle controls. Total RNA was collected from all samples following the respective treatment durations and subjected to RNA sequencing.

### RNA-seq analysis of gene expression changes induced by C-1305 and TRA-01AR

As shown in **Figure 2A** (**Data Set 1**), A549 cells exposed to C-1305 and TRA-01AR for either 24 or 48 hours exhibited similar global gene expression changes, as identified by RNA-seq analysis. However, each compound also elicited distinct and characteristic changes in mRNA expression, indicating quantitative and qualitative differences in the transcriptional responses. The RNA-seq results were normalized to time-matched vehicle-treated controls and interpreted with reference to the transcriptional responses observed in non-cancerous 16HBE14o⁻ cells treated with C-1305 and TRA-01AR under non-cytotoxic conditions. The analysis was then narrowed to the most significant expression changes that were defined as a log2 fold change (FC) ≥ |2| and a false discovery rate (FDR) ≤ 0.05. Importantly, despite both compounds altering the expression of 33 common genes after 24 hours, TRA-01AR uniquely affected 690 mRNAs, while C-1305 influenced 284 mRNAs. After 48 hours, 130 genes were commonly affected, with TRA-01AR additionally altering 623 transcripts and C-1305 affecting 337 others.

**Figure 2.**
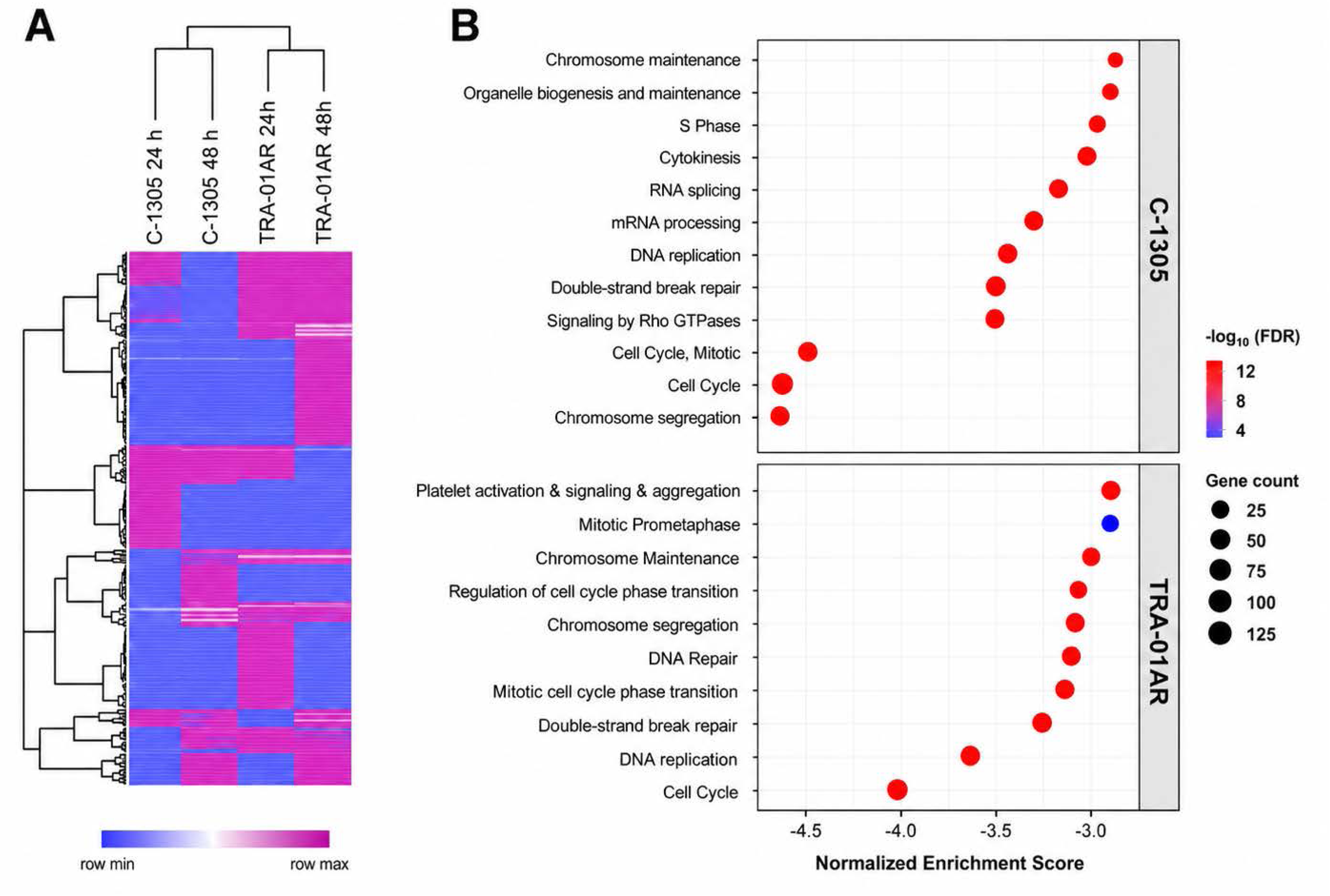
RNA-Iseq analysis of gene expression profiles following treatment with C-I1305 and TRA-I01AR. (**A**) Heatmap of differentially expressed genes in A549 cells treated with C-1305 or TRA-01AR for 24 h and 48 h. Gene expression data were normalized to time-matched vehicle-treated controls. Differentially expressed genes were defined as those exhibiting a log2 fold change (FC) ≥ |2| and a false discovery rate (FDR) ≤ 0.05. Hierarchical clustering was performed using the Morpheus web application. (**B**) Gene Ontology (GO) enrichment analysis of significantly regulated transcripts. Up to ten enriched GO terms are shown for each condition. Enrichment scores were calculated using WebGestalt and are presented as log(p-value) × z-score based on Fisher’s exact test.

Gene ontology (GO) analysis revealed that both compounds, C-1305 and TRA-01AR, significantly impacted pathways associated with DNA replication, DNA repair, and cell cycle regulation, which is consistent with previously reported activities of C-1305, including topoisomerase II inhibition and microtubule dynamics effects (**Figure 2B**). However, distinct biological effects were also observed for each compound. C-1305 upregulated genes involved in RNA processing and Rho GTPase signaling, whereas TRA-01AR downregulated genes associated with platelet activation and signaling pathways. Furthermore, the magnitude of gene expression changes induced by C-1305 was greater than those observed for TRA-01AR.

These findings suggest that while C-1305 and TRA-01AR share a mechanistic overlap, they differ in potency and engage unique molecular pathways that may underlie their distinct cytotoxic profiles.

### Seahorse XF analysis of metabolic parameters following treatment with C-1305 and TRA-01AR

Given the recent evidence linking CRC chemosensitivity to 5-FU with oxidative phosphorylation (OXPHOS) [36], we assessed the impact of C-1305, TRA-01AR, and 5-FU on cellular bioenergetics using the Seahorse XF Analyzer. Real-time measurements of the oxygen consumption rate (OCR) and the extracellular acidification rate (ECAR) were used to determine the mitochondrial respiration and glycolytic flux following the drug exposure. In HCT-116 and HT29 CRC cells (**Figure 3**), the metabolic changes associated with C-1305 and TRA-01AR treatment emerged as early as 6 h, whereas 5-FU-associated bioenergetic alterations became evident primarily at 24 h. The kinetics and magnitude of these shifts differed among compounds and cell lines, indicating both time and cell type-dependent effects.

**Figure 3.**
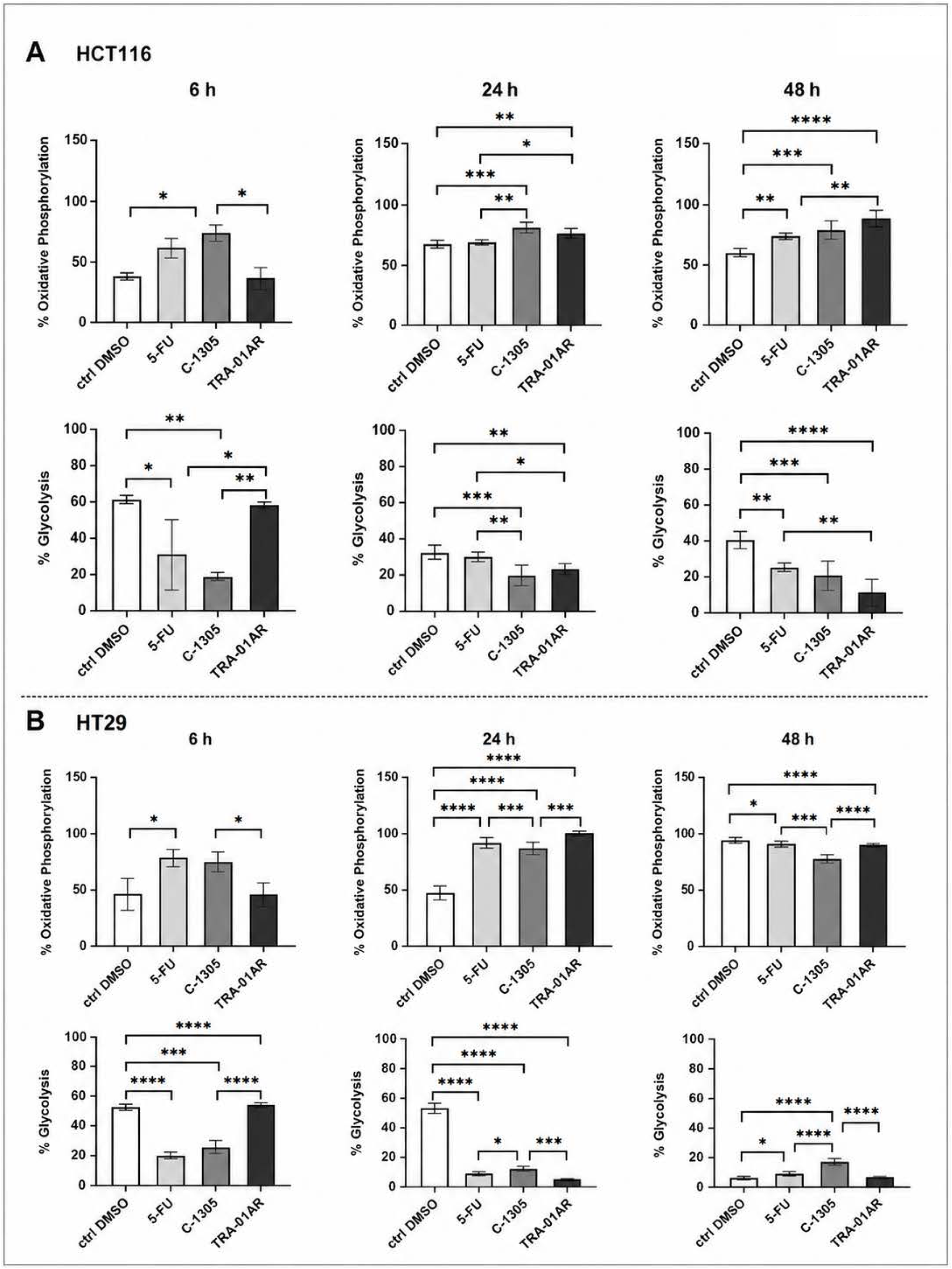
Seahorse XF analysis of metabolic parameters in colorectal cancer cell lines. Real-Time ATP Rate Assay was performed on (**A**) HCT116 and (**B**) HT29 cells following treatment with 5-FU, C-1305, or TRA-01AR for 6 h, 24 h, and 48 h. OCR and ECAR were measured using the Seahorse XF96 Pro Live-Cell system and normalized with Cytation5. Graphs depict the percentage contribution of OXPHOS and glycolysis. CTRL DMSO denotes vehicle-treated controls. Data are presented as mean ± SD with individual values (**Supplemental Table 1**). Statistical analysis was performed using Tukey’s multiple comparisons test*p < 0.05, **p < 0.01, ***p < 0.001, ****p < 0.0001.

In A549 lung cancer cells (**Figure 4B**), treatment with C-1305, TRA-01AR, and 5-FU was associated with changes in the oxygen consumption rate (OCR) and the extracellular acidification rate (ECAR) at both 24 h and 48 h. The magnitude of these changes differed from those observed in non-transformed 16HBE14o⁻ cells (**Figure 4A**).

**Figure 4.**
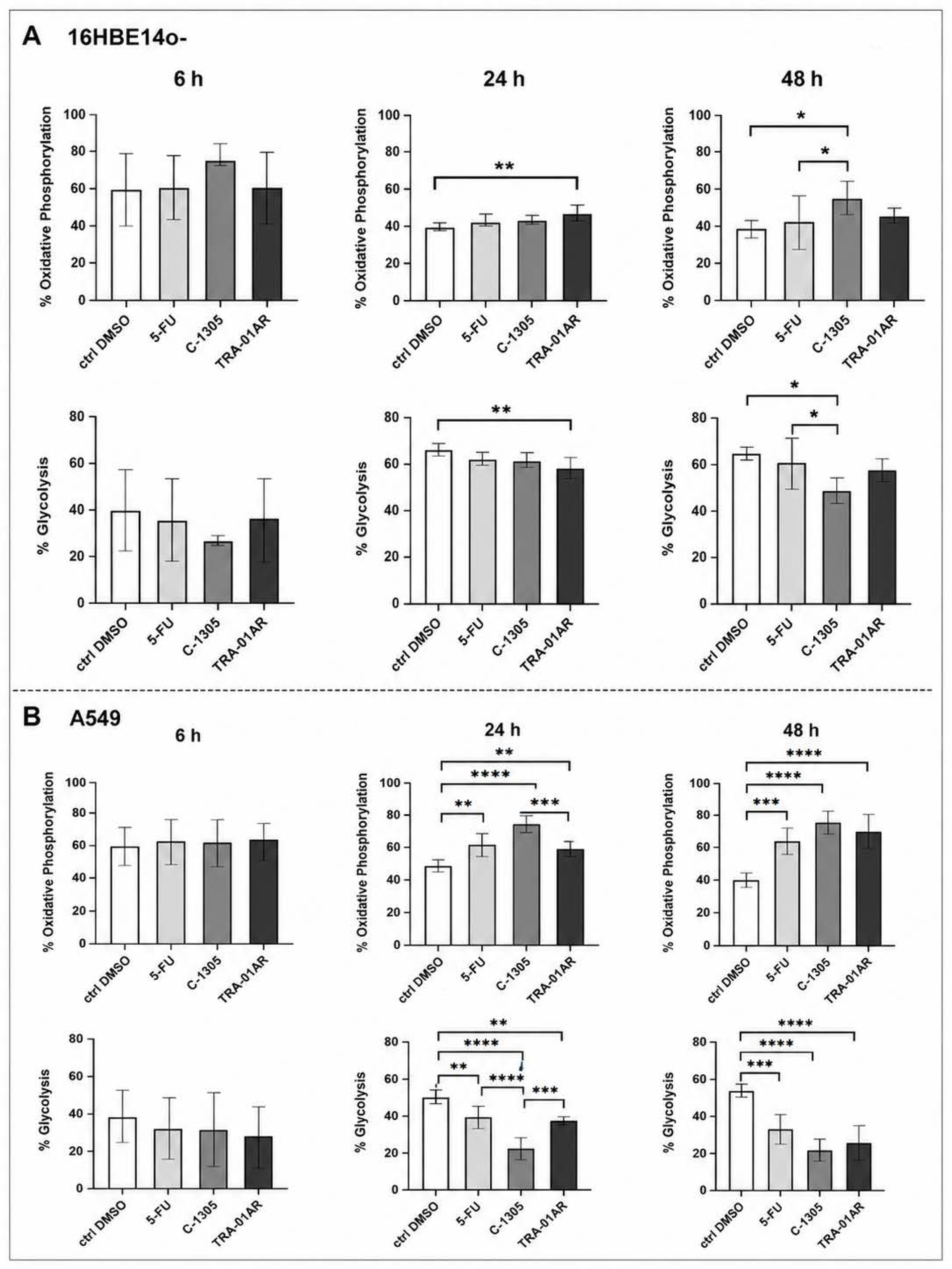
Seahorse XF analysis of metabolic parameters in non-cancerous and cancerous cells following treatment with tested compounds. Real-Time ATP Rate Assay was performed on (**A**) 16HBE14o- and (**B**) A549 cells following treatment with 5-FU, C-1305, or TRA-01AR for 6 h, 24 h, and 48 h. Oxygen consumption rate (OCR) and extracellular acidification rate (ECAR) were measured using the Seahorse XF96 Pro Live-Cell system and normalized with Cytation5 (BioTek, Agilent). Graphs represent the percentage contribution of oxidative phosphorylation (OXPHOS) and glycolysis. CTRL DMSO denotes vehicle-treated controls. Data are presented as mean ± SD with individual values (**Supplemental Table 1**). Statistical analysis was performed using Tukey’s multiple comparisons test*p < 0.05, **p < 0.01, ***p < 0.001, ****p < 0.0001.

In 16HBE14o⁻ bronchial epithelial cells, treatment with TRA-01AR was associated with increased OCR and decreased ECAR at 24 h, whereas similar changes following C-1305 treatment were detected at 48 h. No consistent changes in OCR or ECAR were observed following 5-FU treatment in this cell line within these tested time frames.

Collectively, these data show that C-1305 and TRA-01AR are associated with measurable changes in bioenergetic parameters, occurring earlier and with greater magnitude than those induced by 5-FU, particularly in cancer cells. The observed bioenergetic profiles were broadly consistent with transcriptomic findings, indicating that while C-1305 and TRA-01AR share important mechanistic features, they also induce distinct molecular responses. These observations provided the rationale for evaluating triazoloacridinones in combination with 5-FU *in vivo*. Accordingly, the effects of C-1305 and TRA-01AR were further examined in a CRC xenograft model.

### *In vivo* toxicity assessment of TRA-01AR

The toxicity of TRA-01AR was evaluated using two approaches: modified acute toxicity following a single intravenous (iv) or intragastric (ig) dose, and subchronic toxicity after multiple intravenous administrations that were in accordance with the regulatory guidelines [37–39]. For acute toxicity, female BALB/c mice received TRA-01AR either intravenously or by oral gavage. During a 14-day observation period, no significant signs of toxicity—including changes in body weight, respiration, or behavior—were observed at the highest tested doses of 30 mg/kg (iv) or 250 mg/kg (ig) (**Supplementary Figure 3A)**.

For subchronic toxicity, TRA-01AR was administered intravenously at 30 mg/kg/day for five consecutive days, followed by a 13-day observation period (total 18 days). Only a small weight loss was observed that did not exceed a 10% loss on average (**Supplementary Figure 3B–D**). Notably, previous studies reported that the maximum tolerated dose (MTD) for C-1305 in mice was 150 mg/kg when administered as a single injection or 250 mg/kg when given as daily intraperitoneal injections for 5 days per week [34]. Under the same experimental conditions, TRA-01AR was better tolerated than C-1305.

### RNA-seq analysis of brain, kidney, and liver following C-1305 or TRA-01AR exposure

Global RNA expression profiling was performed on mouse brain, liver, and kidney tissues collected during toxicity assessments after exposure to tolerable doses of C-1305 or TRA-01AR (**Figure 5**; **Data Set 2**). In brain tissue, both compounds reduced expression of genes associated with neuronal systems, neurotransmitter transport, and chemical synapse function. Importantly, these changes, however, were generally less pronounced in the TRA-01AR-treated animals (**Figure 5A**).

**Figure 5.**
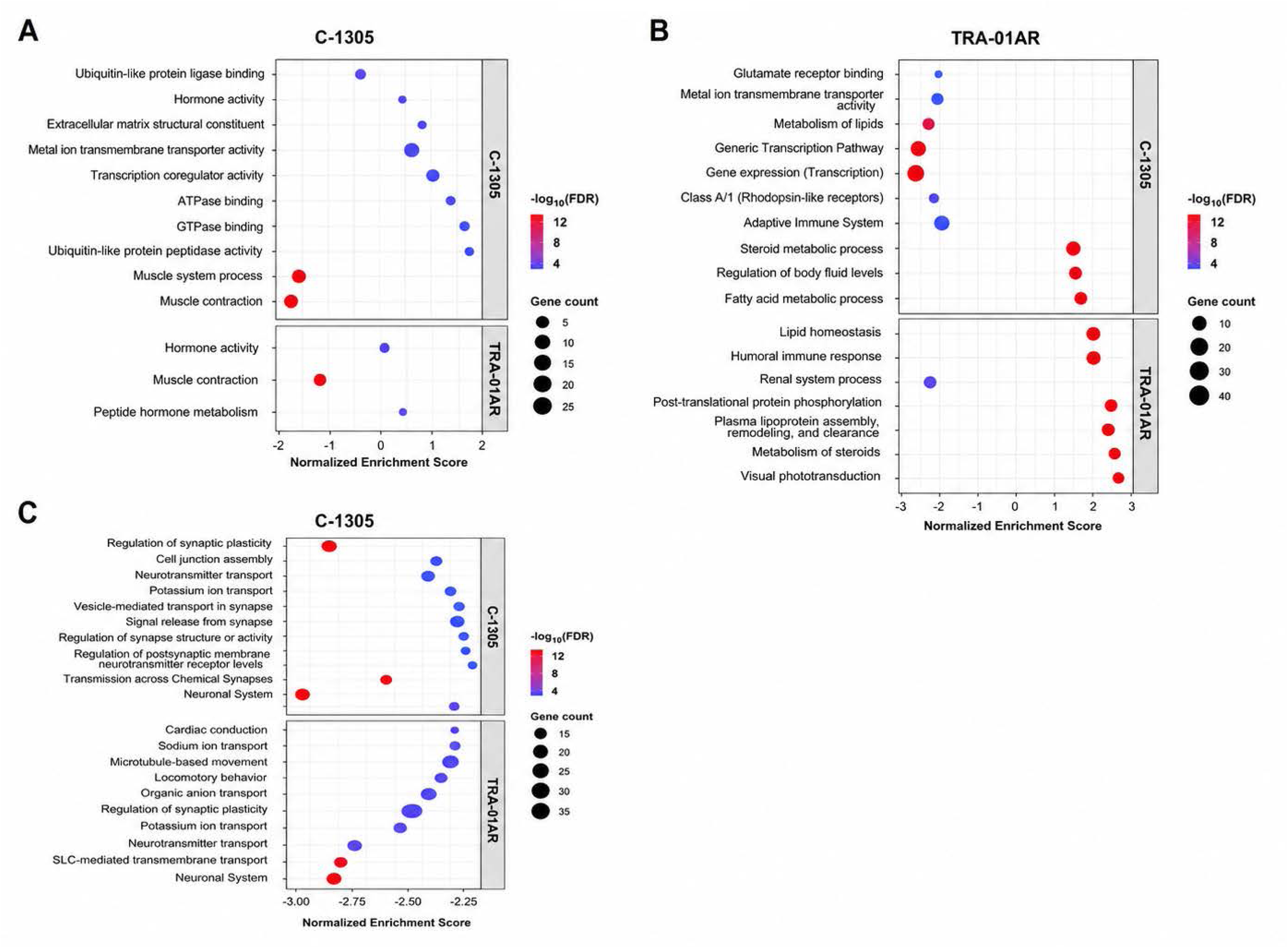
RNA-Iseq analysis of brain, kidney, and liver following treatment with C-I1305 or TRA-I01AR. Gene expression profiles were obtained from brain, kidney, and liver tissues collected during *in vivo* toxicity assessment. RNA from three biological replicates per condition was pooled prior to library preparation and sequencing. Differentially expressed genes were defined as those exhibiting a log2 fold change (FC) ≥ |2| and p < 0.05. (**A**) Brain, (**B**) kidney, and (**C**) liver. Gene Ontology (GO) enrichment analysis was performed using the WebGestalt database, and up to the top 10 enriched GO terms are shown for each tissue. Enrichment scores were calculated from Fisher’s exact test results and are presented as log(p-value) × z-score.

In kidney tissue, the two triazoloacridinones exerted mild but opposing effects on the transcriptomic profiles. C-1305 primarily downregulated genes involved in transcriptional regulation and adaptive immune responses, whereas TRA-01AR modestly upregulated genes linked to body fluid regulation and steroid metabolism while reducing the expression of genes associated with renal system processes (**Figure 5B**).

In liver tissue, TRA-01AR had a minimal impact on genes related to hormone activity and muscle contraction. In contrast, C-1305 induced more substantial changes, and affected genes involved in muscle contraction, hormone activity, extracellular matrix organization, transcriptional regulation, and ATPase/GTPase binding (**Figure 5C**).

Overall, C-1305 treatment elicited a more pronounced transcriptomic response across all examined tissues compared to TRA-01AR, indicating broader systemic transcriptional perturbations under the tested conditions. These findings highlight mechanistic differences between the two compounds and are consistent with the comparatively lower degree of transcriptomic perturbation observed following the TRA-01AR treatment.

### Effects of C-1305 and TRA-01AR in combination with 5-FU in the HT-29 xenograft model

Combination therapy with 5-FU and C-1305 significantly reduced tumor weight in HT-29 xenografts, whereas the 5-FU + TRA-01AR group showed a non-significant trend toward reduced tumor weight (*p* = 0.0863; **Figure 6A**). Tumor perfusion analysis, however, showed increased blood flow parameters in the group treated with 5-FU and TRA-01AR compared to the control conditions (**Figure 6B–C**). The tumor growth kinetics, inhibition rates, and body weight changes are shown in **Supplementary Figure 4**.

**Figure 6.**
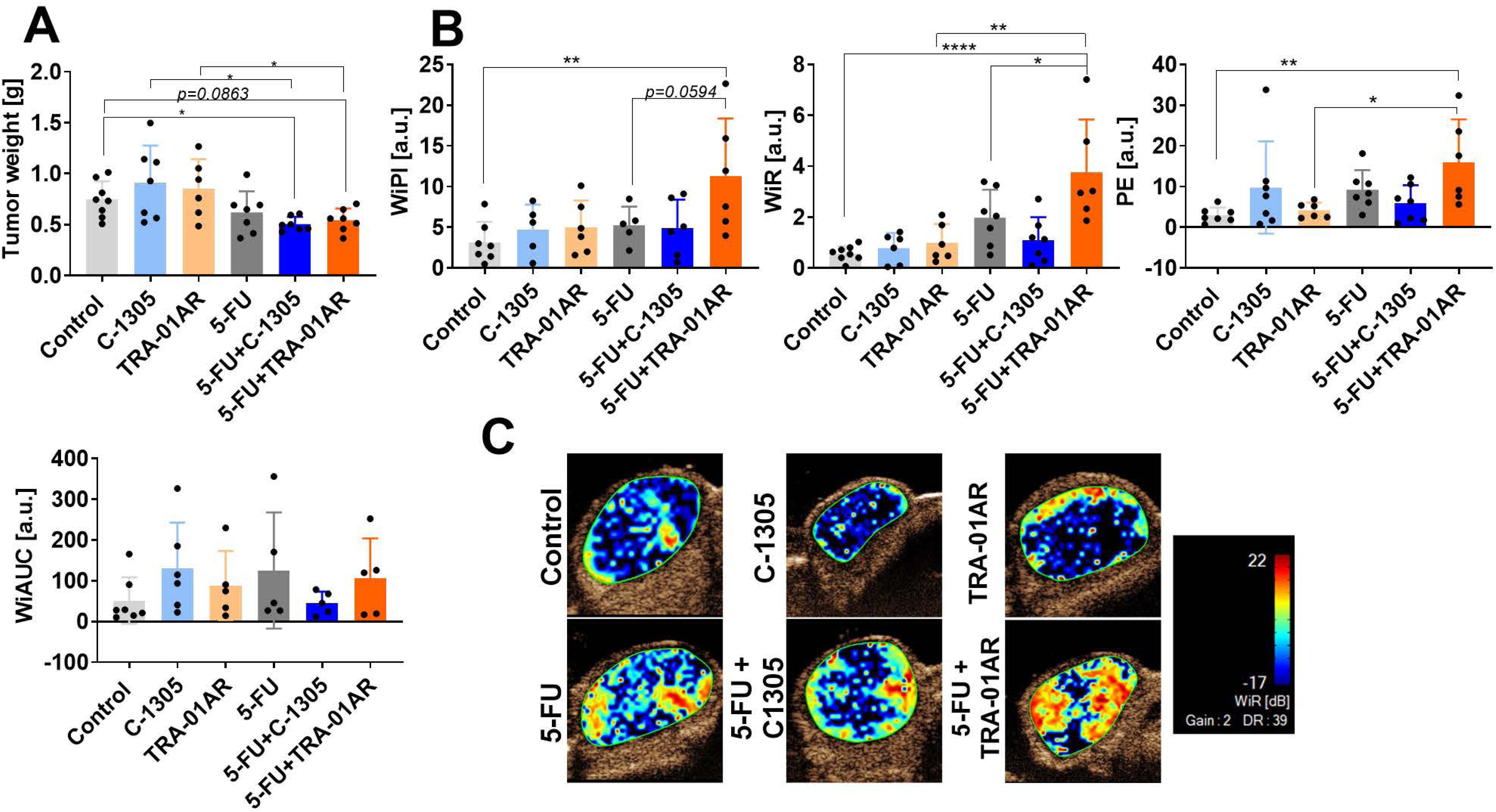
Effects of C-I1305 and TRA-I01AR in combination with 5-IFU in HT-I29 xenografts. (**A**) Tumor weight at endpoint. (**B**) Quantitative analysis of tumor perfusion parameters derived from time-intensity curves (TICs), including peak enhancement (PE), wash-in area under the curve (WiAUC), wash-in rate (WiR), and wash-in perfusion index (WiPI). (**C**) Representative perfusion images (WiR). Data are presented as mean ± SD with individual values. Statistical analysis was performed using Dunnett’s multiple comparisons test; *p < 0.05 vs control.

Histological analysis revealed reduced necrotic areas and decreased CD31 staining in both the combination groups compared to the 5-FU monotherapy group. Given that CD31 is a marker of endothelial cells, these findings suggest treatment-associated alterations in the tumor vascular architecture. Apoptosis (TUNEL assay) was elevated in all treatment groups except in the C-1305 alone group. The TRA-01AR monotherapy upregulated telomerase reverse transcriptase (TERT), an effect that is suppressed by 5-FU in the combination group (**Figure 7A, B**). No major differences were observed in Ki-67, BAX, or α-tubulin. Interestingly, only 5-FU combined with TRA-01AR significantly decreased BAD expression, whereas 5-FU alone increased β-3-tubulin (**Supplementary Figure 5**).

**Figure 7.**
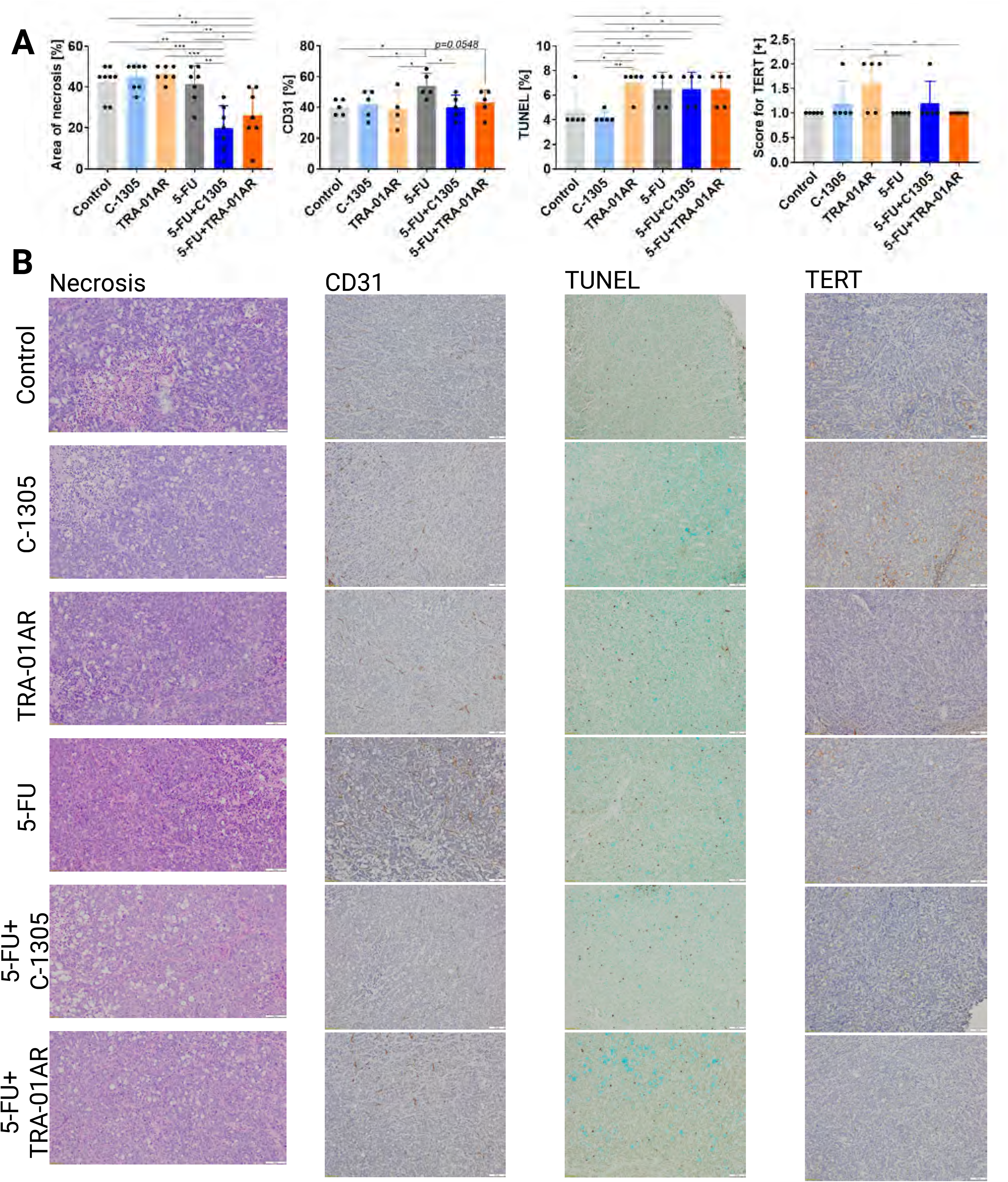
Histological and immunohistochemical analysis of HT-I29 tumors following treatment. (**A**) Quantification of necrotic area, CD31 staining, TUNEL-positive cells, and TERT expression. (**B**) Representative histological and immunohistochemical images corresponding to quantified data. Scale bars indicate 100 µm. Data are presented as mean ± SD with individual values. N= 4-8. Statistical analysis was performed using Dunn’s or Fidher’s LSD (CD31) test; *p < 0.05, **p < 0.01, ***p < 0.001, ****p < 0.0001 vs control.

Plasma cytokine analysis showed elevated interleukin 6 (IL-6) after 5-FU monotherapy, which was reduced by both combinations. C-C motif chemokine ligand 2 (CCL-2) increased with C-1305 (alone and combined with 5-FU) and in the 5-FU + TRA-01AR group (**Figure 8A**). No significant changes in IL-6 or CCL-2 were detected in the tumor tissue (**Figure 8B**).

**Figure 8.**
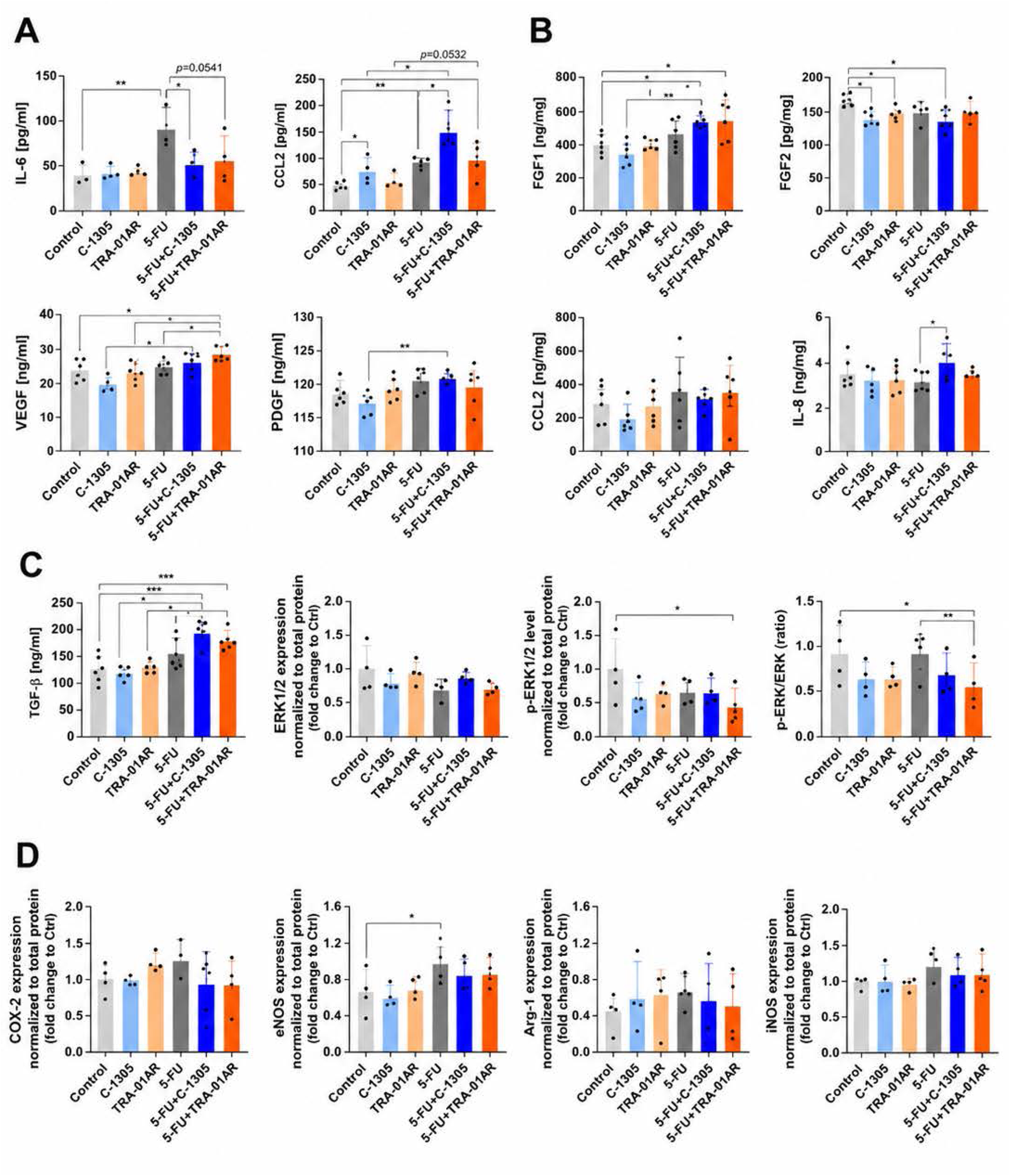
Analysis of plasma and tumor tissue proteins following treatment. (A) Plasma levels of IL-6 and CCL-2 measured by ELISA. (B) Tumor tissue levels of FGF1, FGF2, VEGF, PDGF, CCL-2, IL-6, and TGF-β measured by ELISA. (C) Tumor tissue expression and phosphorylation of ERK1/2 were analyzed using the Jess Simple Western system. (D) Expression of COX-2, eNOS, Ang-4, and p53 were analyzed using the Jess Simple Western system. Data are presented as mean ± SD with individual values. Statistical analysis was performed using Dunnett’s multiple comparisons test and for (D) COX-2, Ang-4 and p-53 Dunn’s multiple comparisons test; *p < 0.05, **p < 0.01, ***p < 0.001, ****p < 0.0001.

In the tumors, fibroblast growth factor 1 (FGF1), vascular endothelial growth factor (VEGF), and Transforming Growth Factor β (TGF-β) were elevated in both combination groups, while fibroblast growth factor 2 (FGF2) decreased with C-1305 and TRA-01AR monotherapy and with C-1305 + 5-FU (**Figure 8B**). The proportion of activated (phosphorylated) extracellular signal-regulated kinase 1 and 2 (p-ERK1/2) compared to the total amount of ERK1/2 protein (p-ERK1/2:ERK1/2 ratio) declined significantly with the 5-FU + C-1305 treatment (**Figure 8C**). No major differences were observed in cyclooxygenase-2 (COX-2), angiogenin 4 (Ang-4), or p53. Only 5-FU monotherapy significantly increased endothelial nitric oxide synthase (eNOS) (**Figure 8D**).

Systemic effects included reduced liver and spleen weight with 5-FU + C-1305 (**Supplementary Figure 6**). Both combinations significantly decreased leukocytes, lymphocytes, and monocytes, while red blood cell count, hemoglobin, and hematocrit were reduced by 5-FU alone and in both combination groups; hemoglobin and hematocrit also decreased with C-1305 monotherapy. Platelet count increased in both combination groups and after C-1305 monotherapy (**Supplementary Figure 7**).

Biochemical analysis revealed unchanged alanine transaminase (ALT) and aspartate transaminase (AST) levels, but the AST/ALT ratio increased with 5-FU + C-1305. Urea levels decreased with TRA-01AR monotherapy and both combinations, whereas creatinine and albumin remained unaffected (**Supplementary Figure 8**).

Taken together, combination therapy with 5-FU and triazoloacridinones reduced tumor growth, but systemic and molecular analyses revealed distinct response profiles. Compared with C-1305, TRA-01AR in combination with 5-FU exhibited a distinct systemic and molecular profile and a comparatively reduced impact on selected systemic parameters under the tested conditions.

## DISCUSSION

5-Fluorouracil (5-FU) remains a cornerstone of CRC treatment, yet its clinical efficacy is limited by resistance, toxicity, and incomplete tumor control. These challenges highlight the need for combination strategies that enhance the therapeutic response while maintaining tolerability. In this study, we applied a multidisciplinary approach integrating chemical modification, cellular and transcriptomic profiling, bioenergetic analysis, and *in vivo* validation to evaluate triazoloacridinones as potential modulators of the 5-FU response.

A key conceptual advance of this work is the identification of bioenergetic modulation as an additional and previously underexplored dimension of triazoloacridinone activity. Both C-1305 and TRA-01AR induced a shift toward oxidative phosphorylation accompanied by reduced glycolytic activity. This observation is consistent with emerging evidence linking metabolic reprogramming, particularly increased mitochondrial dependency, to reduced responsiveness to fluoropyrimidines in CRC [36, 40, 41]. Specifically, 5-FU-resistant CRC cells have been shown to exhibit increased dependence on OXPHOS to sustain survival and stem-like properties under treatment pressure [41]. In this context, mitochondrial metabolic reprogramming has emerged as a key adaptive mechanism contributing to disease progression and therapeutic resistance in CRC [40]. Within this framework, our findings suggest that triazoloacridinones may alter the metabolic context in which 5-FU exerts its effects. However, this interpretation remains hypothesis-generating, given that no direct causal relationship between metabolic changes and therapeutic responses was established in this study.

At the transcriptional level, both compounds affected pathways associated with DNA replication, repair, and cell cycle regulation, consistent with the known activity of C-1305 as a topoisomerase II-targeting agent with additional effects on cytoskeletal dynamics [22, 26]. Notably, C-1305 induced broader and more pronounced transcriptional changes than TRA-01AR, which may explain C-1305’s higher apparent cytotoxic potency but also its less favorable systemic effects *in vivo*. In contrast, TRA-01AR exhibited more moderate and context-dependent transcriptional responses, suggesting that chemical modification can decouple potency from systemic impact.

The BOC modification introduced in TRA-01AR was designed to alter physicochemical properties and intracellular engagement. Prodrug strategies are widely employed to improve physicochemical properties and enable controlled activation of pharmacologically active compounds following intracellular or tissue-specific processing [42]. The delayed cellular responses observed for TRA-01AR compared with C-1305 are consistent with altered intracellular dynamics; however, direct evidence for prodrug-like activation or intracellular conversion is lacking. No pharmacokinetic or metabolomic analyses were performed to assess compound processing or activation. Therefore, any interpretation of TRA-01AR as a prodrug should be considered provisional and requires further validation.

The *in vivo* data further illustrate the functional differences between the two compounds. While C-1305 in combination with 5-FU significantly reduced tumor burden, TRA-01AR in combination produced only a non-significant trend toward tumor reduction and did not improve efficacy relative to 5-FU alone. These findings suggest that the bioenergetic and transcriptional effects observed *in vitro* are not sufficient in their current form to translate into enhanced antitumor activity *in vivo*. Importantly, TRA-01AR displayed a more favorable systemic profile, suggesting a trade-off between efficacy and tolerability, which may be relevant for future optimization of dosing strategies and treatment schedules.

In addition, TRA-01AR uniquely increased intratumoral blood flow, pointing to a potential effect on tumor vascular function. Tumor perfusion and microvascular dynamics are increasingly recognized as critical determinants of an effective therapeutic response, influencing drug delivery and intratumoral distribution in CRC [43]. Although improved perfusion may enhance drug penetration, the functional consequences of this effect remain unclear, as no corresponding improvement in tumor control was observed. Therefore, vascular modulation should be interpreted cautiously and not considered as evidence for any therapeutic enhancement.

The present study follows a two-step experimental strategy in which mechanistic insights were first obtained in a well-characterized epithelial model (A549), followed by validation in CRC systems. This approach enables high-resolution analysis of cellular responses but also introduces important limitations related to model specificity. The use of A549 cells for mechanistic analyses is supported by our previous work demonstrating their suitability for identifying key molecular targets of C-1305, including microtubule stabilization and modulation of the IRE1α/XBP1 pathway [22, 25]. However, such findings require validation in disease-specific contexts, particularly in CRC. Importantly, previous transcriptomic analyses performed in HT-29 cells treated with C-1305 revealed partially overlapping gene expression signatures, supporting the relevance of mechanistic observations derived from A549 cells in CRC-derived systems [22, 25, 28]. Together, these findings support the use of A549 cells as a robust platform for mechanistic profiling, while highlighting the need for validation in CRC models.

Several limitations should be considered when interpreting these findings. First, the mechanistic investigations, including transcriptomic analyses, were primarily performed in a non-CRC model. Second, the functional validation of key transcriptomic and metabolic observations remains limited. Third, the proposed mechanisms, including the role of bioenergetic reprogramming, vascular effects, and the potential intracellular activation, remain correlative rather than causal. Finally, the absence of pharmacokinetic and metabolic characterization limits the conclusions regarding compound processing and activity.

Despite these limitations, this study provides a novel integrative framework linking chemical modification to coordinated transcriptional, metabolic, and physiological effects. Rather than establishing immediate therapeutic applicability, the findings define key mechanistic hypotheses and highlight bioenergetic modulation [44] as a potential axis of interaction with fluoropyrimidine-based therapy.

Taken together, our results indicate that triazoloacridinones influence multiple cellular programs relevant to cancer progression and treatment response. However, further studies are required to establish their mechanistic basis, validate their activity in CRC-specific systems, and determine whether these effects can be translated into a meaningful therapeutic benefit.

## CONCLUSION

Triazoloacridinones modulated the cellular and molecular processes relevant to the tumor response, including bioenergetic, transcriptional, and signaling-related pathways. Distinct differences between C-1305 and TRA-01AR were observed across experimental systems, reflecting divergence in potency, systemic impact, and biological effects.

*In vivo*, the combination of 5-FU with C-1305 resulted in a significant reduction in tumor weight, whereas the combination with TRA-01AR did not achieve a statistically significant antitumor effect compared to 5-FU alone. Notably, TRA-01AR was associated with increased intratumoral blood flow and a comparatively reduced impact on systemic parameters under the tested conditions, suggesting a distinct balance between biological activity and tolerability.

These findings indicate that triazoloacridinones can influence cellular programs relevant to therapy response; however, they do not establish a direct therapeutic advantage of TRA-01AR in combination with 5-FU under the conditions tested. Instead, the observed effects support further investigation to clarify its mechanism of action, optimize dosing strategies, and evaluate its potential relevance in CRC–specific contexts.

## METHODS

### Compound synthesis and characterization

#### Compound C-1305

(C_18_H_19_N_5_O_2_, M_W_ = 337.38 g/mol., InChI=1S/C18H19N5O2/c1-22(2)9-3-8-19-13-5-6-14-17-16(13)18(25)12-10-11(24)4-7-15(12)23(17)21-20-14/h4-7,10,19,24H,3,8-9H2,1-2H3) was synthesized in several steps by adopting an optimization of the literature descriptions [6,10]. In the first step, 6-chloro-3-nitro-2-{[4-(benzyloxy)-phenyl]amino}benzoic acid was obtained by Ullman condensation of 2,6-dichloro-3-nitrobenzoic acid and 4-(benzyloxy)aniline hydrochloride in the presence of triethylamine in anhydrous ethanol at reflux. The next step was cyclization of the above anthranilic acid derivative by the action of phosphorus oxychloride at reflux. Subsequent stages of the synthesis were the removal of the benzyl group from the 8-OH group, reduction of 4-nitroacridin-9(10H)-one derivative to the respective 4-aminoacridin-9(10H)-9-one, followed by diazotization of the 4-amino group. The last step of the C-1305 synthesis route was the substitution of the halogen atom (5-Cl) in the latter compound by 3-(dimethylamino)propan-1-amine. Purification of C-1305 was performed by applying gravitational liquid chromatography using silica gel (SiO_2_, Merck) as a stationary phase and a gradient type of elution (chloroform/methanol/ammonia mixtures). Detailed information concerning the synthesis of compound C-1305 and its isolation in chemically pure form is provided in [22]. The compound purity and identity were verified with RP-HPLC, elemental analysis, ^1^H NMR spectroscopy, and HR MS spectrometry. Representative chromatograms and spectra together with experimental settings are presented in [22]. The C-1305 chemical formula is provided in **Figure 1A**. The two independent batches of C-1305 were tested. C-1305 was stored in the dark at 4 °C. Prior to the experiments, the compound was dissolved in DMSO in a 3 mM stock solution.

#### To obtain TRA-01AR

(C_23_H_27_N_5_O_4_, MW = 437.49 g/mol, InChI = 1S/C23H27N5O4/c1-23(2,3)32-22(30)31-14-7-10-18-15(13-14)21(29)19-16(24-11-6-12-27(4)5)8-9-17-20(19)28(18)26-25-17/h7-10,13,24H,6,11-12H2,1-5H3). Purified C-1305 (20 mg) was dissolved in dichloromethane (CH₂Cl₂). A five-fold molar excess of triethylamine (41 μL) and di-tert-butyl dicarbonate (62 μL) was added, and the reaction mixture was stirred at room temperature for 2 h. Upon completion, the solvent and residual triethylamine were removed under reduced pressure. The crude product was purified by flash chromatography on neutral alumina (Al₂O₃) using CHCl₃/hexane (1:1, v/v) as the mobile phase since preliminary studies demonstrated the chemical instability of the compound on silica gel. The purified product was obtained as a yellow solid powder (yield: ∼20 mg, ∼69%). The reaction scheme is shown in **Figure 1D**.

The obtained compound was characterized by high-resolution mass spectrometry (HRMS), reverse-phase high-performance liquid chromatography (RP-HPLC), melting point analysis, and proton nuclear magnetic resonance (¹H NMR) spectroscopy. HRMS analysis revealed a molecular ion peak at m/z 438.2148, consistent with the calculated value of 438.2261. Chemical purity was assessed by RP-HPLC using a Shimadzu LC-20AD system equipped with a photodiode array detector (SPD-M20A) and a Supelco Discovery C8 analytical column (250 × 4.6 mm, 5 μm). Elution was performed using a linear gradient of solvent A (80% acetonitrile in water containing 0.08% trifluoroacetic acid) and solvent B (100% acetonitrile) over 60 min at a flow rate of 1 mL/min. Chromatograms were monitored in the 190–800 nm range. TRA-01AR exhibited a chromatographic purity of 94.52%. The ¹H NMR spectrum recorded in CD₃OD displayed characteristic signals corresponding to the acridone core, methylene linker, and *tert*-butyl moiety, with signal assignments supported by COSY correlations. The melting point was determined using a Büchi M-565 apparatus. Collectively, these analyses support the structural assignment and analytical characterization of TRA-01AR (**Supplementary Figure 1**).

For biological studies, TRA-01AR was converted to its hydrochloride salt. Briefly, a slight molar excess of hydrogen chloride in anhydrous diethyl ether was added dropwise at 5°C to a solution of TRA-01AR in a methanol/chloroform mixture. The resulting yellow precipitate was collected and recrystallized from an ethyl ether/methanol mixture. Two independent batches of TRA-01AR were used throughout the study. The compound was stored in the dark at 4°C and dissolved in DMSO to a stock concentration of 3 mM prior to use. DMSO (catalog no. S-002-D) was purchased from Sigma-Aldrich.

### Cell lines and culture conditions

Experiments were performed using a panel of human colon cancer cell lines (HT-29, HCT-116, DLD-1, LoVo, LS180, LS1034, SW-480, and SW-620) as well as the normal intestinal epithelial line FHC. In addition, human A549 lung adenocarcinoma cells and human lung microvascular endothelial cells (HLMEC) were included in selected experiments. All experiments were performed in at least three independent biological replicates unless otherwise stated. The cell lines used in this study are listed in Supplementary Table S1, together with their origin, culture conditions, and reference catalog numbers.

**Supplementary Table 1.**
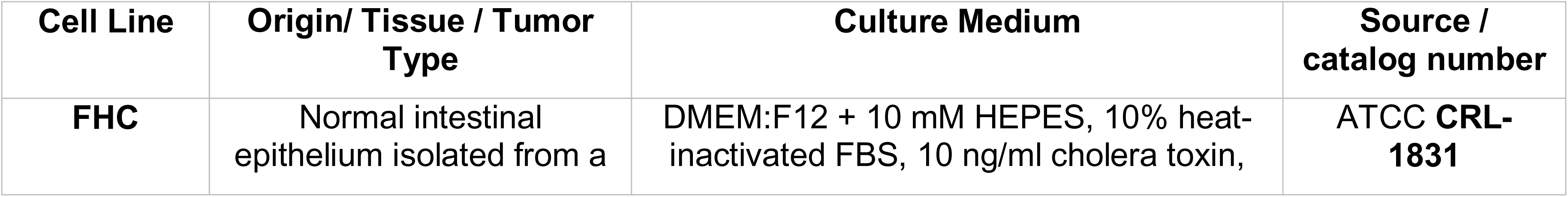
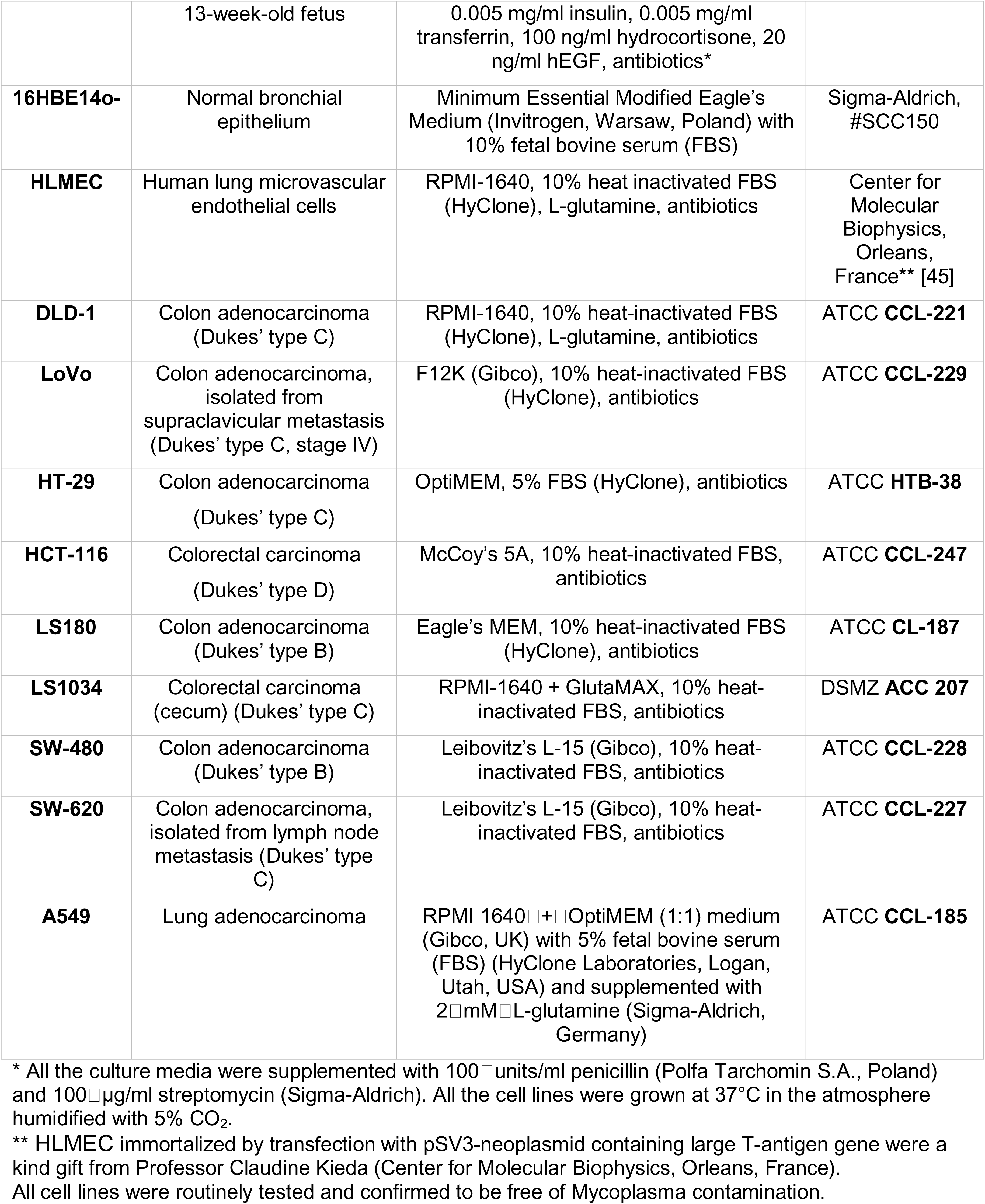
Characteristics of human cancer and normal cell lines used in this study.

| Cell Line | Origin/ Tissue / Tumor Type | Culture Medium | Source / catalog number |
|---|---|---|---|
| **FHC** | Normal intestinal epithelium isolated from a 13-week-old fetus | DMEM:F12 + 10 mM HEPES, 10% heat-inactivated FBS, 10 ng/ml cholera toxin, 0.005 mg/ml insulin, 0.005 mg/ml transferrin, 100 ng/ml hydrocortisone, 20 ng/ml hEGF, antibiotics* | ATCC **CRL-1831** |
| **16HBE14o-** | Normal bronchial epithelium | Minimum Essential Modified Eagle's Medium (Invitrogen, Warsaw, Poland) with 10% fetal bovine serum (FBS) | Sigma-Aldrich, #SCC150 |
| **HLMEC** | Human lung microvascular endothelial cells | RPMI-1640, 10% heat inactivated FBS (HyClone), L-glutamine, antibiotics | Center for Molecular Biophysics, Orleans, France** [45] |
| **DLD-1** | Colon adenocarcinoma (Dukes' type C) | RPMI-1640, 10% heat-inactivated FBS (HyClone), L-glutamine, antibiotics | ATCC **CCL-221** |
| **LoVo** | Colon adenocarcinoma, isolated from supraclavicular metastasis (Dukes' type C, stage IV) | F12K (Gibco), 10% heat-inactivated FBS (HyClone), antibiotics | ATCC **CCL-229** |
| **HT-29** | Colon adenocarcinoma (Dukes' type C) | OptiMEM, 5% FBS (HyClone), antibiotics | ATCC **HTB-38** |
| **HCT-116** | Colorectal carcinoma (Dukes' type D) | McCoy's 5A, 10% heat-inactivated FBS, antibiotics | ATCC **CCL-247** |
| **LS180** | Colon adenocarcinoma (Dukes' type B) | Eagle's MEM, 10% heat-inactivated FBS (HyClone), antibiotics | ATCC **CL-187** |
| **LS1034** | Colorectal carcinoma (cecum) (Dukes' type C) | RPMI-1640 + GlutaMAX, 10% heat-inactivated FBS, antibiotics | DSMZ **ACC 207** |
| **SW-480** | Colon adenocarcinoma (Dukes' type B) | Leibovitz's L-15 (Gibco), 10% heat-inactivated FBS, antibiotics | ATCC **CCL-228** |
| **SW-620** | Colon adenocarcinoma, isolated from lymph node metastasis (Dukes' type C) | Leibovitz's L-15 (Gibco), 10% heat-inactivated FBS, antibiotics | ATCC **CCL-227** |
| **A549** | Lung adenocarcinoma | RPMI 1640□+□OptiMEM (1:1) medium (Gibco, UK) with 5% fetal bovine serum (FBS) (HyClone Laboratories, Logan, Utah, USA) and supplemented with 2□mM□L-glutamine (Sigma-Aldrich, Germany) | ATCC **CCL-185** |

\* All the culture media were supplemented with 100□units/ml penicillin (Polfa Tarchomin S.A., Poland) and 100□µg/ml streptomycin (Sigma-Aldrich). All the cell lines were grown at 37°C in the atmosphere humidified with 5% CO<sub>2</sub>.
\*\* HLMEC immortalized by transfection with pSV3-neoplasmid containing large T-antigen gene were a kind gift from Professor Claudine Kieda (Center for Molecular Biophysics, Orleans, France).
All cell lines were routinely tested and confirmed to be free of Mycoplasma contamination.

### Cytotoxicity assays

MTT assays of cell viability were performed in 96-well plates [46]. One day after plating, 16HBE14o− cells (15,000 cells per well), A549 cells (7,500 cells per well) or HCT-116 cells (2,000 cells per well) were cultured with different concentrations of compounds. Cell line–specific seeding densities were optimized experimentally based on proliferation rates to ensure comparable growth conditions and to avoid overconfluence during the assay. Under vehicle-treated control conditions, all cell lines reached approximately 70–80% confluency at the time of analysis. After a 24-hour incubation with the specified compounds, medium containing 1 mg/mL MTT was added to cells for a final concentration of 0.5 mg/mL and incubated at 37 °C for 4 h. The medium was aspirated, and the formazan product was solubilized with DMSO. The absorbance at 630 nm (background absorbance) was subtracted from absorbance at 570 nm for each well. Six technical replicates were used for each concentration within each independent experiment. DMSO was used as control and the concentration in the medium did not exceed 0.1%. The IC_50_ values were calculated from dose–response curves by nonlinear regression using the four-parameter logistic function. All IC_50_ values were derived from at least three independent biological experiments.

Sulforhodamine B (SRB) assay was conducted in 96-well plates (Greiner Bio-One, Kremsmünster, Austria). After 24 h of culture under standard conditions, C-1305 and TRA-01AR were added at concentrations ranging from 0.1 to 100 µg/mL, and 5-FU at concentrations ranging from 0.01 to 10 µg/mL or DMSO (0.1%, solvent control), and incubated for 72 h. The SRB protocol was carried out using a BioTek dosing and rinsing station (BioTek Instruments, Inc., Winooski, VT, USA). Cells were fixed for 1 h with 25% trichloroacetic acid at room temperature (RT), then washed three times with tap water. Subsequently, cells were stained with 0.1% sulforhodamine B (prepared as a w/v solution in 1% acetic acid) for 30 min. Next, plates were rinsed four times with 1% acetic acid and protein-bound dye was extracted with 10 mM unbuffered Tris base for 30 min. Absorbance was measured at a wavelength of λ_=_540 nm using a Synergy H4 hybrid microplate reader (BioTek Instruments, Inc., Winooski, VT, USA). The IC50 values were calculated from dose–response curves using the four-parameter logistic function.

### Isolation of RNA

Total RNA, including the small non-coding RNA fraction, was isolated using the miRNeasy Kit (Qiagen, Cat. No. 217004) according to the manufacturer’s protocol. RNA concentration and purity were determined spectrophotometrically based on absorbance at 260 and 280 nm. Samples were stored at −70_°C until use. RNA integrity was assessed using the Agilent TapeStation system.

### Next Generation RNA sequencing analyses (*in vitro*)

Cells treated with C-1305, TRA-01AR or DMSO vehicle (as specified in the Results section) were used for the RNA isolation and analyses. Following total RNA isolation, samples were validated with quantitative real-time PCR for apoptosis activation prior to further analysis. Following rRNA depletion, the remaining RNA fraction was used for library construction and subjected to 75-bp single-end sequencing on an Illumina NextSeq 550 instrument (San Diego, CA, USA). Sequencing reads were aligned to the Gencode human reference genome assembly (GRCh38 p13 Release 32) using STAR version 2.7.3a [47] (using parameters --outReadsUnmapped Fastx --outSAMtype BAM SortedByCoordinate --outSAMattributes All). Transcript assembly and estimation of the relative abundance and tests for differential expression were carried out with HTSeq-count version 0.11.3 (using parameters -m union -r pos -t exon -i gene_id -a 10 -s no -f bam) and DESeq2 (using default parameters) [48–50]. Selected RNA-seq findings were independently assessed by quantitative real-time PCR. WebGestalt [51] was used to place the Next Generation Sequencing (NGS) results into a physiological context. Furthermore, the analyses were limited to experimentally verified interactions and no extended gene enrichment set analyses were performed. The results of WebGestalt analysis were further verified with Enrichr [52] as we did in [53]. Gene ontology (GO) visualizations were generated using SRplot [54].

### Seahorse XF metabolic assays

Each cell line, 16HBE14o-, A549, HCT116, and HT29, was cultured under standardized conditions in their respective media as described above. Following the treatments with DMSO (negative control), 5-FU, C1301, or TRA-01AR at specific time points of 6, 24, or 48 hours, the culture media was replaced with DMEM XF Assay Media (Agilent Technologies, Santa Clara, CA). This media was freshly supplemented prior to the experiment with glucose (25 mM), pyruvate (1 mM), and L-glutamine (2 mM). Before each assay, cells underwent a 1-hour incubation for degassing, followed by brightfield imaging using the Cytation5 Cell Imaging Multimode Reader (BioTek, Agilent, USA). During the ATP Rate Assay (Agilent Seahorse ATP Rate Assay Kit), the oxygen consumption rate (OCR) and extracellular acidification rate (ECAR) were measured before and after sequential injections of 1.5 µM oligomycin and 1 µM rotenone/antimycin A mix using the Seahorse XF96 Pro Analyzer. For each experiment, four to six technical replicates were seeded per cell line. Normalization was performed using fluorescence Hoechst staining on the Cytation5 reader and expressed as specific result values per 1,000 cells. Data analysis was conducted using the dedicated Seahorse Wave software (Agilent). Current Seahorse XF methodologies require cell seeding in specialized microplates, which can be challenging for certain primary or differentiated epithelial cell types that are not readily adaptable to standard assay formats [55]. Accordingly, normal colon-derived epithelial cells were not included in Seahorse analyses due to technical limitations. Preliminary experiments yielded low and highly variable oxygen consumption rates, resulting in an insufficient signal-to-noise ratio for reliable quantification and interpretation. This observation is consistent with recognized limitations of extracellular flux assays in primary epithelial systems [55].

### In vivo toxicity assessment

Experiments were carried out on 8-to 10-week-old female and male BALB/c mice (purchased from the Tri-City Academic Laboratory Animal Centre — Research and Services Centre of the Medical University of Gdansk, Poland). Female mice were used to test toxicity of the new compound – TRA-01AR. Toxicity of C-1305 had already been established in previous studies. The principles of the 3Rs (Replacement, Reduction, and Refinement) allowed us to utilize male mice that were otherwise scheduled for euthanasia, thereby avoiding the need to use additional animals while still confirming the compound’s toxicity profile. For the acute toxicity study, one mouse for each dose tested was used during the initial phase and 4 mice were used to confirm the Maximum Tolerated Dose in the main phase. For subchronic toxicity, 5 mice were used for each dose. Approval of the study was obtained from the Local Ethical Committee for Animal Experiments in Bydgoszcz (permission number: 19/2018). All the experiments were performed according to Directive 2010/63/EU of the European Parliament and Council on the protection of laboratory animals used for scientific purposes. Mice were bred and housed in the Tri-City Academic Laboratory Animal Centre — Research and Services Centre of the Medical University of Gdansk, Poland. Animals were maintained on a regular, 12-h light-dark cycle at 22 ± 2°C and provided with freely available food and water.

Toxicity assessment was performed using a modified Fixed-Dose method for both acute and subchronic toxicity studies. This method, intended to reduce animal use, involved sequential dosing aimed at finding the dose that causes apparent toxicity but not death [56]. An acute toxicity study was carried out in two separate phases: an initial phase to identify the Maximum Tolerated Dose (MTD) and a main phase to confirm it. During the initial phase, mice (n = 1 per tested dose, male for C-1305 and female for TRA-01AR) were injected intravenously via the tail vein at increasing doses of TRA-01AR ranging from 5 to 50.0 mg/kg in a 5% glucose solution for injection, or the tested compound was administered by gavage with doses from 5 to 250 mg/kg in a mixture of 5% DMSO and 95% PBS (doses from 5 to 100 mg/kg), or 5% glucose for injection for higher doses. Doses were given until clear but non-fatal toxic symptoms appeared. Signs such as body weight change, coat condition, respiration, cage environment, behavior, and mobility were evaluated pre-administration every 15 minutes during the first hour post-administration, and once daily thereafter for 14 days. After observation, the animals were humanely euthanized, and internal organs were collected for further analysis. Brains, liver and kidney tissue samples of control animals as well as mice injected intravenously with TRA-01AR at a dose of 30 mg/kg or C-1305 at a dose of 20 mg/kg were collected for follow-up transcriptomic analysis. This approach was consistent with the Fixed-Dose Procedure recommendations intended to minimize animal use [56].

During the main phase of acute toxicity assessment, only one dose, identified as the MTD in the initial phase, was tested for each route of administration. TRA-01AR was injected intravenously at a dose of 30 mg/kg (n = 4) or administered via oral gavage at a dose of 250 mg/kg (n = 4). Mice were observed before and after administration and euthanized according to the same protocol described for the initial phase. Subchronic toxicity was evaluated solely for intravenous administration. A dose of 30 mg/kg/day of TRA-01AR in a 5% glucose for injection solution was given (n = 5) over five consecutive days. Changes in body weight, coat condition, respiration, cage environment, behavior, and mobility were assessed before administration, and every 15 minutes during the first hour post-administration, throughout the administration period, and once daily for 13 days following the final dose. After observations, the animals were humanely euthanized.

### Next Generation RNA sequencing analyses (in vivo tissues)

Kidney, brain, and liver tissues were collected from mice and immediately snap-frozen in liquid nitrogen. Approximately 10 mg of each tissue sample was stored at −80_°C until further processing. For RNA isolation, tissues were homogenized in QIAzol Lysis Reagent (Qiagen) following the manufacturer’s protocol. Total RNA was extracted using the standard phenol-chloroform method and purified according to Qiagen guidelines. RNA quality and integrity were assessed using spectrophotometric analysis (NanoDrop) and electrophoretic profiling (TapeStation) to ensure RNA Integrity Number (RIN) ≥ 9. After successful isolation, three biological replicates per condition for each tissue type were pooled in equal proportions (1:1:1) to generate representative samples. This approach was adopted to obtain representative tissue-level transcriptional profiles while minimizing sequencing costs. This illustrates that the resulting analyses should be considered exploratory and hypothesis-generating. Pooled RNA after rRNA depletion was then used for library preparation and subjected to 75-bp single-end sequencing on an Illumina HiSeq 2000 instrument (San Diego, CA, USA). Sequencing reads were aligned to the Gencode mouse reference genome assembly (GRCm38 p6 Release M24) using STAR version 2.7.3a [47]. Transcript assembly and estimation of the relative abundance and tests for differential expression were carried out with HTSeq-count version 0.11.3 with Cufflinks and Cuffdiff [48–50]. The resulting data were validated with quantitative real-time PCR. WebGestalt [51] was used to place the Next Generation Sequencing results into a physiological context. Furthermore, the analyses were limited to experimentally verified interactions and no extended gene enrichment set analyses were performed. The results of WebGestalt analysis were further verified with Enrichr [52] as we did in [53]. Gene ontology visualization has been performed with SRplot [54].

### Quantitative Real-Time PCR

Total RNA was extracted from 50 mg of snap-frozen liver using the Epicenter MasterPure RNA purification kit (MC85200, Biozym), in accordance with the manufacturer’s instructions. cDNA was reverse-transcribed using an RT^2^ First Stand Kit (330404; Qiagen, Germany). Real-time quantitative PCR was performed using the AbiPrism 7500 sequence detection system and Taqman fast advanced master mix (4444556, Applied Biosystems, Germany) with the following assay probes: Gapdh (Mm_99999915_g1), Ctsd (Mm00515586_1), Ctsb (Mm01310506_m1), Tfeb (Mm00448968_m1), TFEB (Hs01065085_m1), and 18S (Hs03003631_m1). All probes were purchased from Life Technologies (Carlsbad, CA, USA). Gene expression levels were quantified and analyzed in technical duplicates. The mRNA expression levels of the target genes were compared to controls, normalized against the reference gene Gapdh or 18S rRNA and quantified using the 2^−ΔΔCt method.

#### HT-29 xenograft model and treatment protocol

Experiments were carried out on 6-to 8-week-old female BALB/c nude mice (CByJ.Cg-Foxn1^nu^/J purchased from Charles River Laboratories). Each test group consisted of 8 mice. Approval for the study was obtained from the Local Ethical Committee for Animal Experiments in Wroclaw (permission number: 012/2020). All the experiments were performed according to Directive 2010/63/EU of the European Parliament and Council on the protection of laboratory animals used for scientific purposes. Animals were housed under specific pathogen-free conditions and exposed to 12-h day/night cycles with access to feed and water ad libitum at the animal facility of Hirszfeld Institute of Immunology and Experimental Therapy in Wroclaw, Poland.

Mice were injected subcutaneously on the right flank with HT-29 cells suspended in 0.15 ml phosphate-buffered saline (3 × 10_ cells) mixed with 0.05 ml Matrigel (3:1, v/v; Matrigel Basement Membrane Matrix, High Concentration). Treatment was initiated when tumors reached ∼130 mm³ (day 10 after cell inoculation). C-1305 was dissolved in 5% glucose and administered intraperitoneally (30 mg/kg) on days 10, 11, 12, 13, 24, 25, and 26. TRA-01AR (dissolved in 5% glucose) was administered intraperitoneally at 20 mg/kg on the same schedule. 5-Fluorouracil (5-FU, Ebewe Pharma; 50 mg/ml) was given intravenously at 50 mg/kg once per week for 3 weeks (day 10, 17, and 24). Animals were monitored daily and euthanized after 28 days. Body weight and tumor size were measured three times per week. Tumor volume (TV) was calculated as TV (mm³) = 0.5 × a × b², where *a* is the largest diameter and *b* the perpendicular diameter. Blood and tumor tissue were collected at autopsy for further analyses.

### Tumor perfusion analysis

On day 18, tumor perfusion was assessed using the Vevo 2100 ultrasound system (VisualSonics) with the MicroMarker™ contrast agent. The agent was dissolved in 1 ml sterile saline, and ∼100 μl was injected intravenously. Mice were anesthetized with 2–3% isoflurane in synthetic air (600 ml/min) and positioned on the imaging table. Tumors were covered with bubble-free gel and visualized in transverse cross-section using the MS250 scanhead. Imaging included a bolus sequence (∼15 fps) after contrast signal stabilization (∼50 s), followed by burst-mode destruction of microbubbles and a replenishment sequence. After imaging, mice recovered in a warm environment. Data were analyzed with Vevo LAB 1.7.1 and VevoCQ software (VisualSonics).

### Toxicity profile of the anticancer treatment

The body weight of the HT-29 tumor-bearing animals was measured thrice each week throughout the course of all the studies. Whole blood parameters were analyzed using a Mythic 18 hematology analyzer (PZ Cormay, Warsaw, Poland). Biochemical analyses were performed in Cobas C 111 analyzer (Roche Diagnostics) for the molecules alanine aminotransferase (ALTL), aspartate aminotransferase (ASTL), urea (UREL), creatinine (CRE2) and albumin (ALB) using reagents and procedures provided by the manufacturer.

### Histological and immunohistochemical analyses

HT-29 tumors were frozen in liquid nitrogen and stored at −80 °C. Approximately 300 mg of frozen tissue was homogenized in 500 μl RIPA buffer supplemented with protease and phosphatase inhibitors (Sigma-Aldrich) using the FastPrep®-24 system (2 cycles, 6 m/s, 40 s). Homogenates were incubated on ice for 20 min, frozen, thawed, and centrifuged (4 °C, 15 min, 14,000 g). Supernatants were collected, centrifuged again, and stored at −80 °C. Protein concentration was determined by the modified Lowry assay (Bio-Rad).

HT-29 tumors were analyzed using standard methods at Sorbolab Research Laboratory LLC, Poznań, Poland. Standard staining techniques with haematoxylin and eosin and TUNEL assay were performed. Moreover, tumor slides were stained with anti-Ki67, anti-CD31, anti-telomerase reverse transcriptase (TERT), anti-α-tubulin and anti-β-3-tubulin, anti-BAD, and anti-BAX antibodies (all from Thermo Fisher Scientific), counterstained with haematoxylin, and finally observed under Leica DM 200 microscope. The results were analyzed using an immunoreactive score (IRS) based on the percentage of stained cells and staining intensity.

### Protein and cytokine analyses (Western blot / ELISA)

Tumor lysates were analyzed using the Jess Simple Western System (ProteinSimple) for phosphorylation and expression of Ang-4, COX-2, p53, ERK1/2, phospho-ERK1/2, and eNOS (R&D Systems or CST antibodies). Lysates were diluted to 2 µg/µL (COX-2, p53, ERK1/2, phospho-ERK1/2) or 0.5 µg/µL (eNOS, Ang4), mixed with sample buffer and master mix, and denatured (95 °C, 5 min). Samples (5 µl) and biotinylated ladders were loaded, and proteins were detected by chemiluminescence or NIR fluorescence. Total protein was quantified in the same run using biotin-labelling and streptavidin-HRP. Data were analyzed with Compass software (v6.0.0; ProteinSimple) and normalized to total protein. Results are expressed as fold change relative to controls.

### Detection of proteins in mice plasma using ELISA

The plasma samples of mice injected with HT-29 tumor cells were analyzed for the expression levels of CCL2 and IL-6 proteins by using enzyme-linked immunosorbent assay (ELISA; Invitrogen) according to the manufacturer’s protocol. For this purpose, the blood samples were collected in EDTAK2 tubes and centrifuged at 2400×_*g* for 14_min at 4°C. Absorbance was recorded at 450_nm using a BioTek Synergy H4 hybrid reader (Biokom).

### Detection of proteins in tumor lysates using ELISA

The lysates obtained from HT-29 tumors were analyzed for the expression levels of VEGF, PDGF, CCL2, FGF1, FGF2, TGF-β, and IL-6 proteins by using ELISA (EIAab Science) according to the manufacturer’s protocol. Absorbance was recorded at 450_nm using a BioTek’s Synergy H4 hybrid reader (Biokom). Results were normalized to total protein concentration in each lysate.

### Statistical analysis

Data are presented as mean ± SD or as indicated in the figure legends. Data distribution was assessed using the Shapiro–Wilk normality test. Depending on the experimental design and data distribution, appropriate statistical tests were applied, followed by post hoc multiple-comparison tests where applicable. Statistical analyses were performed using GraphPad Prism version 10.6.0 (GraphPad Software, San Diego, CA, USA). A p-value < 0.05 was considered statistically significant. Details of the statistical analyses performed for individual datasets, including test statistics and p-values, are provided in the corresponding figure legends and **Supplementary Table S1**.

## DECLARATIONS

## Materials availability

Further information and requests for resources and reagents may be directed to and will be fulfilled by the corresponding authors.

## Data and code availability

All data reported in this paper will be shared by the lead contact upon request. This paper does not report original code. Any additional information required to reanalyze the data reported in this paper is available from the lead contact upon request.

## Acknowledgments

This research was funded by This work was funded by National Center for Research and Development Program under contract Strategmed3/306853/9/NCBR/2017 (TARGETTELO) to M.B.; R.B.; J.W; M.M. and by DFG SFB 1177 (259130777, F11), to L.S.

## Author contributions

Marta Świtalska and Beata Filip-Psurska performed animal experiments with tumor-bearing mice. Magdalena Milczarek analyzed protein expression in tumor tissue. Mateusz Psurski conducted tumor blood flow analyses. Eliza Turlej and Magdalena Milczarek tested the antiproliferative activity of the compounds. Damian Szczesny and Michał Markuszewski performed animal toxicity experiments. Sylwia Bartoszewska and Magda Gebert carried out MTT assays and RNA isolation for transcriptomic analysis. Paulina Czechowicz performed Seahorse XF bioenergetic analysis. Kamil Krzymiński and Małgorzata Gawrońska synthesized, characterized, and provided the compounds. David Crossman and Katarzyna Wierzbicka performed bioinformatic analysis of NGS data., Maciej Bagiński, Leszek Kalinowski, and Liliana Scheafer contributed to the discussion. Joanna Wietrzyk participated in the *in vivo* experiments, analyzed the results, and contributed to manuscript writing. Rafal Bartoszewski participated in transcriptomic experiments, analyzed the results, and wrote the original manuscript draft. James F. Collawn revised the manuscript.

## Declaration of interests

The authors declare that they have no competing interests.

## Declaration of generative AI and AI-assisted technologies

Generative AI language tools (Microsoft Copilot) were used to assist in correcting language and grammar in this document. No content was generated beyond linguistic refinement.

**Supplementary Figure 1.** Physicochemical character-ation of TRA-01AR. The melting point of TRA-01AR was determined using a Büchi M-565 apparatus according to the manufacturer’s protocol. Compound purity was assessed by reverse-phase high-performance liquid chromatography (RP-HPLC), demonstrating a chromatographic purity of 94.52%. Representative RP-HPLC chromatograms are shown. Compound identity was verified by high-resolution electrospray ionization quadrupole time-of-flight mass spectrometry (ESI-QTOF), which revealed a molecular ion peak at m/z 438.2148, consistent with the calculated value of 438.2261. Structural assignment was further supported by ¹H NMR spectroscopy recorded in CD₃OD at room temperature, displaying characteristic signals corresponding to the aromatic acridone core, methylene linker, and *tert*-butyl moiety. Signal assignments were validated by COSY correlations.

**Supplementary Figure 2.** Half-maximal inhibitory concentration (IC₅₀) of the tested compounds in several colon cancer cell lines, normal hepatocytes, and endothelial cells. Cells were incubated with compounds for 72 h, and the SRB assay was performed. Data were analyzed using D. Nevozhay’s Cheburator software. N=3-4 with three technical replicates in each biological replicate. Data are presented as mean ± SD. Statistical analysis: two-way ANOVA revealed a significant interaction between cell line and compound (F(18,59) = 12.83, p < 0.0001), indicating that the effects of the compounds differed among cell lines. Significant main effects of cell line (F(9,59) = 22.25, p < 0.0001) and compound (F(2,59) = 81.37, p < 0.0001) were also observed. Šídák’s multiple comparisons test: *p < 0.05, **p < 0.01, ***p < 0.001, ****p < 0.0001 compared with 5-FU for each cell line.

**Supplementary Figure 3.** Toxicological assessment of TRA-01AR. (**A**) Body weight of mice treated with a single dose of TRA-01AR at the maximum tolerated dose (MTD) via intravenous and intragastric routes. N=4. (**B**–**E**) Subchronic toxicity evaluation of TRA-01AR administered in five daily doses of 30 mg/kg (total cumulative dose: 150 mg/kg). N=5. (**B**) Mean body weight ± SD and (**C**) body weight of individual animals. (**D**) Mean body weight change (%) ± SD and (**E**) body weight change in individual animals. Arrows indicate the days of compound administration.

**Supplementary Figure 4.** Tumor growth and body weight of mice bearing HT-29 human colon cancer and treated with 5-Fu, C-1305, TRA-01AR alone or combined. (A) Kinetics tumor growth measured during the experiment; (B) tumor growth inhibition (TGI, percent); (C) kinetics of body weight measured during the experiment; (D) body weight change (BWC, percent). Data are presented as (A and C) 25th to 75th percentiles (box) and min to max (whiskers) with individual values or (B and D) mean percent of TGI or BWC. N=6-8 (**Supplemental Table 1**). Statistical analysis: Fisher’s LSD test; *p < 0.05, **p < 0.01, ***p < 0.001, ****p < 0.0001.

**Supplementary Figure 5.** Immunohistochemical analyses of HT-29 tumor tissue harvested from mice treated with 5-Fu, C-1305, TRA-01AR alone or combined. Expression of (A) Ki-67, (B) BAD, (C) BAX, (D) α-tubulin, and (E) β-3-tubulin. Representative immunohistochemical images corresponding to quantified data. Scale bars indicate 100 µm. Data are presented as mean ± SD with individual values. N=4-5 (**Supplemental Table 1**).. Statistical analysis was performed using Dunn’s test; *p < 0.05 vs control.

**Supplementary Figure 6.** Weight of organs from mice bearing HT-29 tumors and treated with 5-Fu, C-1305, TRA-01AR alone or combined. (A) Liver, (B) kidneys, (C) spleen weights. Data are presented as mean ± SD with individual values. N= 6-8 (**Supplemental Table 1**).. Statistical analysis was performed using (A and C) Tukey’s multiple comparisons test or (B) Dunn’s multiple comparisons test; *p < 0.05, **p < 0.01, ***p < 0.001, ****p < 0.0001.

**Supplementary Figure 7.** Morphological parameters of blood from mice bearing HT-29 tumors and treated with 5-Fu, C-1305, TRA-01AR alone or combined. (A) Count for white blood cells (WBC), (B) distribution of lymphocytes, monocytes and granulocytes, (C) red blood cells (RBC) parameters including RBC count, hemoglobin concentration and hematocrit, (D) platelets count. Left panel: data for C-1305 alone or combined; right panel: data for TRA-01AR alone or combined. Data are presented as 25th to 75th percentiles (box) and min to max (whiskers) with individual values. N= 6-8 (**Supplemental Table 1**). Statistical analysis was performed using Fisher’s LSD test; *p < 0.05, **p < 0.01, ***p < 0.001, ****p < 0.0001.

**Supplementary Figure 8.** Biochemical parameters of plasma from mice bearing HT-29 tumors and treated with 5-Fu, C-1305, TRA-01AR alone or combined. (A) ALT, alanine aminotransferase (B) AST, aspartate aminotransferase; (C) AST/ALT, ratio of aspartate to alanine aminotransferase; (D) CRE, creatinine; (E) UREA, urea; (F) ALB, albumin. Data are presented as mean ± SD with individual values. N= 6-8 (**Supplemental Table 1**). Statistical analysis was performed using (A and E) Dunn’s test or (B-D and F) Fisher’s LSD test; *p < 0.05, **p < 0.01, ***p < 0.001, ****p < 0.0001.

**Data Set 1. Transcriptomic results of TRA01-AR and C-1305 treatments of human cells *in vitro* along with GO assignments.**

**Data Set 2. Transcriptomic results of TRA01-AR and C-1305 treatments of human cells *in vivo* along with GO assignments.**

## Abbreviations

5FU: 5-fluorouracil
18S: 18S ribosomal RNA
ALB: albumin
ALT: alanine aminotransferase
Ang4: angiogenin-4
ANOVA: analysis of variance
AST: aspartate aminotransferase
BAD: Bcl-2-associated death promoter
BAX: BCL2-associated X protein
*CCL2*: C-C motif chemokine ligand 2
CD31: cluster of differentiation 31
COX-2: cyclooxygenase-2
CRC: colorectal cancer
CST: Cell Signaling Technology antibodies
*CTSB*: cathepsin B
*CTSD*: cathepsin D
DMEM: F12: Dulbecco’s Modified Eagle Medium:Ham’s F-12 Nutrient Mixture
DMSO: dimethyl sulfoxide
ECAR: extracellular acidification rate
EGFR: epidermal growth factor receptor
eNOS: endothelial nitric oxide synthase
ERK1/2: extracellular signal-regulated kinase 1/2
ESI-QTOF: electrospray ionization quadrupole time-of-flight
FBS: fetal bovine serum
FGF1/2: fibroblast growth factor 1/2
FHC: fetal human colon cells
*GAPDH*: glyceraldehyde 3-phosphate dehydrogenase
GO: gene ontology
HEPES: 4-(2-hydroxyethyl)-1-piperazineethanesulfonic acid
HLMEC: human lung microvascular endothelial cells
HRMS: high-resolution mass spectrometry
HRP: horseradish peroxidase
HTSeq-count: High-throughput sequencing read counting
*IL-6*: interleukin 6
MEM: Minimum Essential Medium
MTD: maximum tolerated dose
MTT: 3-(4,5-dimethylthiazol-2-yl)-2,5-diphenyltetrazolium bromide
NGS: Next Generation Sequencing
NMR: nuclear magnetic resonance
OCR: oxygen consumption rate
OXPHOS: oxidative phosphorylation
p53: tumor protein p53
PBS: phosphate-buffered saline
PDGF: platelet-derived growth factor
PE: peak enhancement
qRT-PCR: Quantitative real-time polymerase chain reaction
RIN: RNA Integrity Number
RIPA: Radio Immuno Precipitation Assay buffer
RPMI-1640: Roswell Park Memorial Institute medium
rRNA: ribosomal RNA
RT: rise time
SRB: sulforhodamine B
STAR: Spliced Transcripts Alignment to a Reference
TERT: telomerase reverse transcriptase
*TFEB*: transcription factor EB
*TGFβ1*: Transforming growth factor beta 1
TICs: time–intensity curves
TS: thymidylate synthase
TUNEL: terminal deoxynucleotidyl transferase dUTP nick end labeling
VEGF: vascular endothelial growth factor
WiAUC: wash-in area under the TIC curve
WiPI: wash-in perfusion index
WiR: wash-in rate or maximum slope

## Notes

### Competing Interest Statement

The authors have declared no competing interest.

