## Supplementary Figures for "Preclinical Evaluation of a Novel BOC-Modified Triazoloacridinone (TRA-01AR) in Combination with 5-Fluorouracil in Colorectal Cancer"

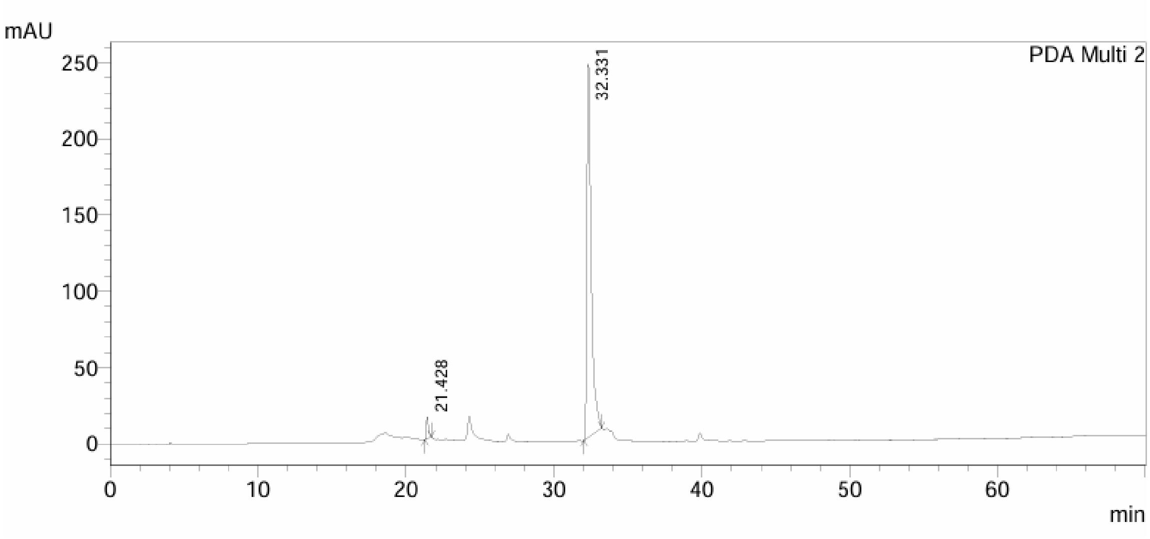


RP-HPLC chromatograms of TRA-01AR (UV detection at 254 nm). The chemical purity of the TRA-01AR was over 94.52% and was determined using high-performance liquid chromatography (HPLC) qualitative analyses (LC-20AD, Shimadzu) on reverse-phase column (250 × 4.6 mm; Supelco Discovery Analytical column) which containing C-8 material (5 µm particle size). TRA-01AR was analyzed with a linear gradient of the mixture of the two solvent systems A and B in 60 min (0-100% B) at a flow rate of 1 ml/min, where solvent A consisted of 80% acetonitrile in water acidified with 0.08% trifluoroacetic acid, whereas solvent B was 100% acetonitrile. The analysis was verified using a photodiode array detector (SPD-M20A; Shimadzu) over the range, 190 to 800 nm.

#### Supplementary Figure 1

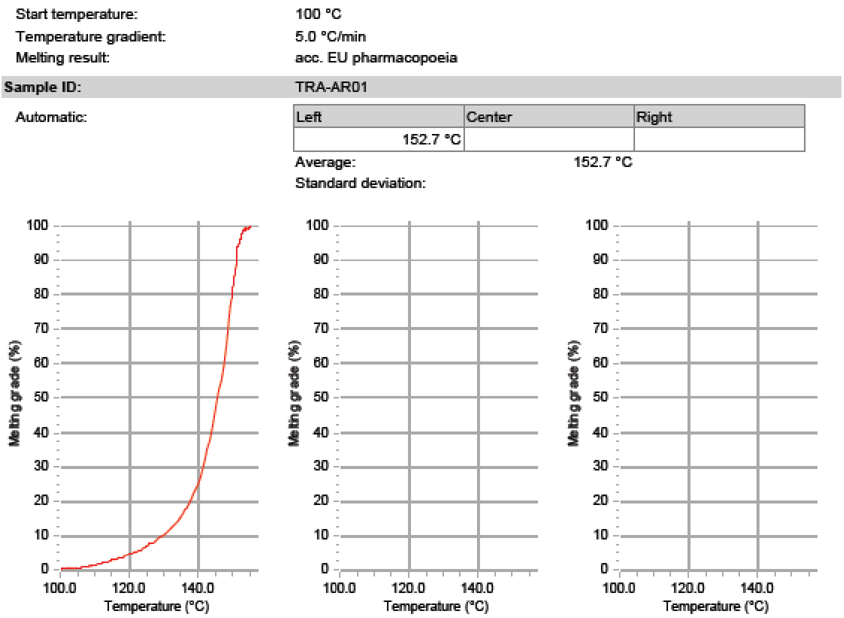


The melting point of TRA01-AR was measured using a Büchi M-565 melting point apparatus according to the manufacturer's instructions. The sample was placed in a sealed capillary tube, and the temperature was gradually increased until complete melting was observed. The melting point range was recorded.

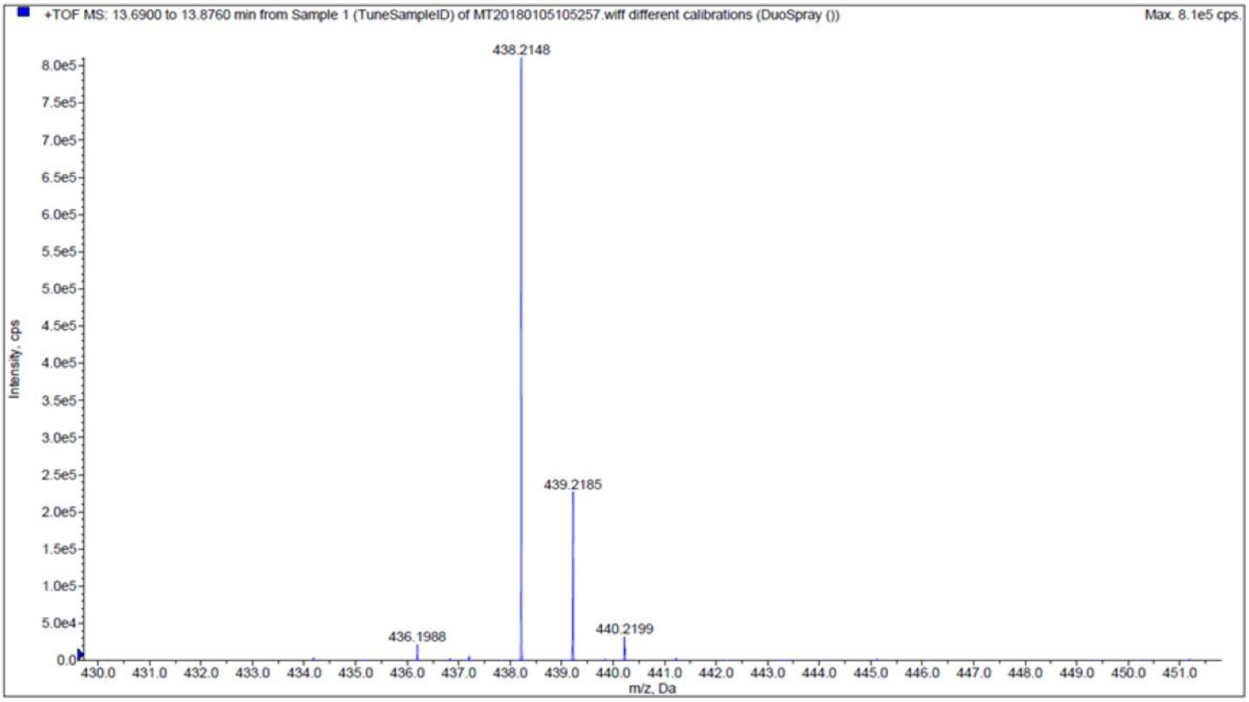


The identity of TRA01-AR was confirmed by electrospray ionization quadrupole time-of-flight (ESI-QTOF) mass spectrometry. The spectrum showed a prominent peak corresponding to the molecular ion ($M^+$) with an m/z value of 438.2148, which is consistent with the theoretical value of 438.2261. No significant fragment ions or impurities were detected, confirming the purity of the compound.

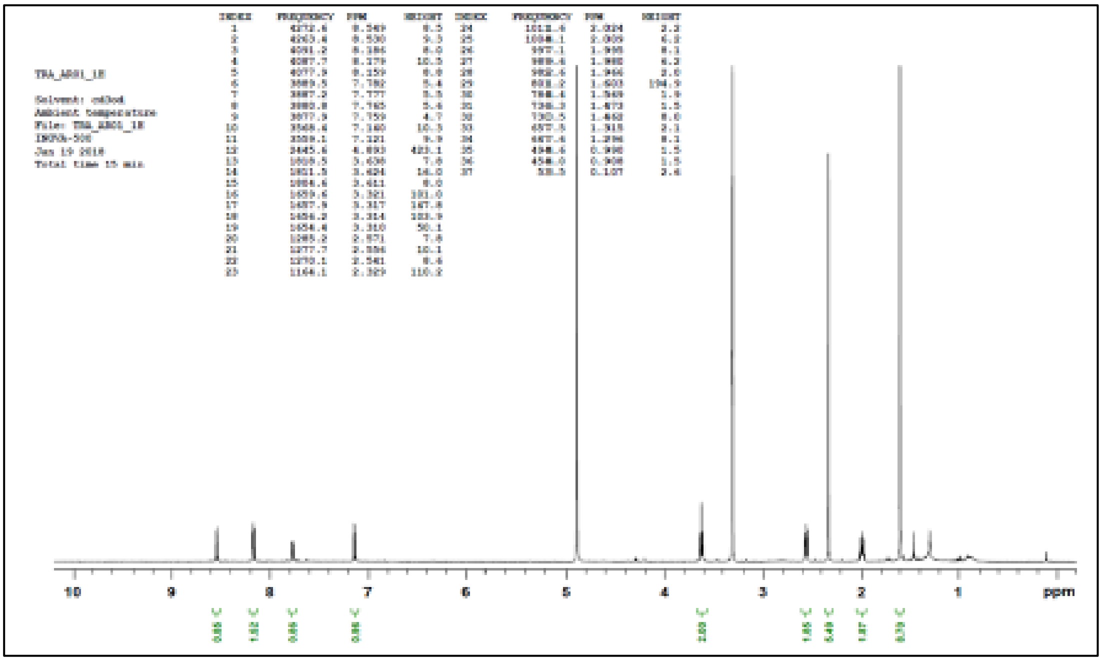


| TRA01-AR | $^1$H NMR chemical shifts ($CD_3OD$) |
|---|---|
| 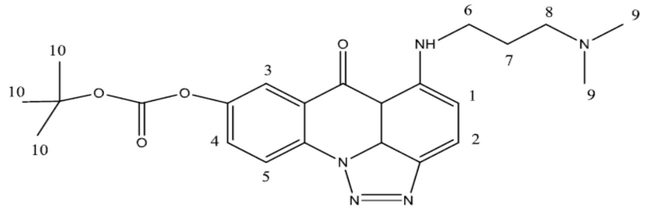 | 1. δ 7.782, 7.777, 7.765, 7.759 |
| | 2. δ 8.549, 8.530 |
| | 3. δ 8.179 |
| | 4. δ 8.186, 8.159 |
| | 5. δ 7.121, 7.140 |
| | 6. δ 3.638, 3.624, 3.611 |
| | 7. δ 2.024, 2.009, 1.995, 1.980, 1.966 |
| | 8. δ 2.571, 2.556, 2.541 |
| | 9. δ 2.329 |
| | 10. δ 1.603 |

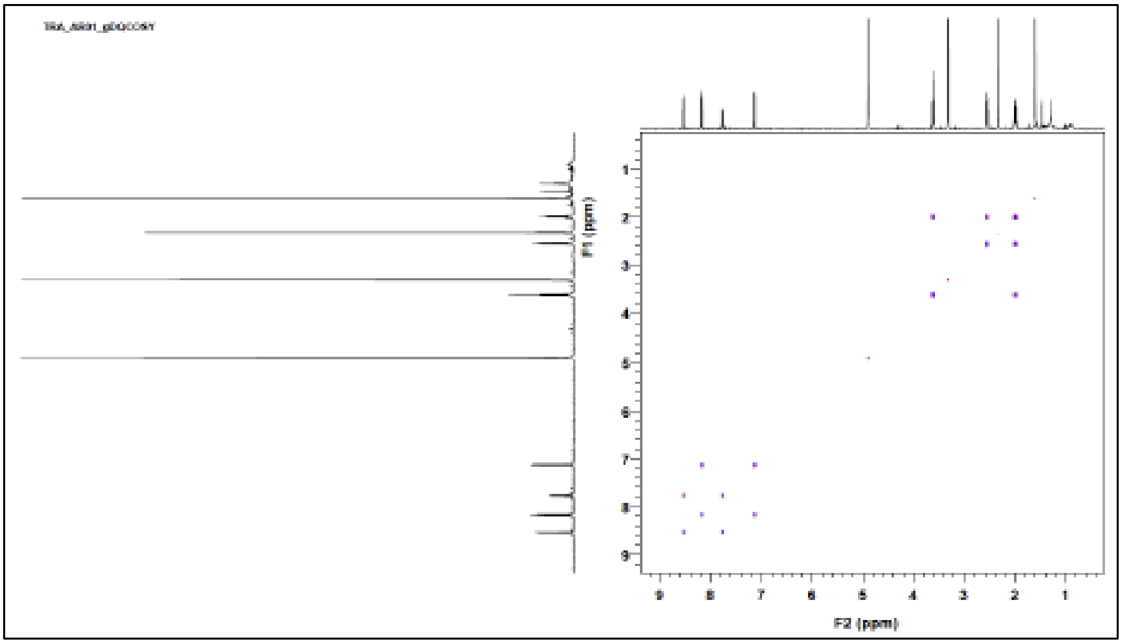


The structure of TRA01-AR was confirmed by $^1$H NMR spectroscopy recorded in $CD_3OD$ at room temperature. The spectrum showed characteristic signals corresponding to aromatic protons of the acridone core, methylene groups of the side chain, and tert-butyl groups from the BOC protecting group. Coupling patterns were verified using COSY correlations.

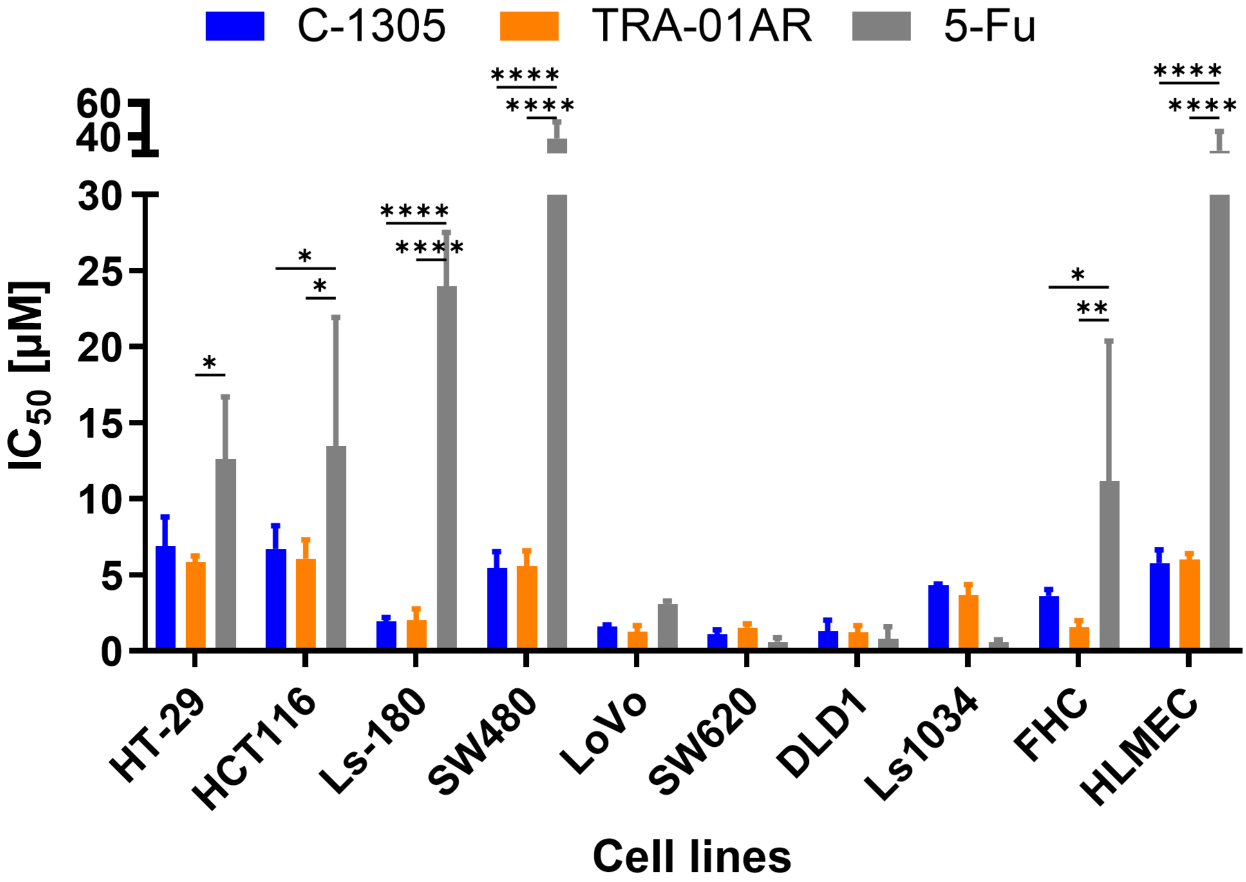
C-1305
TRA-01AR
5-Fu
$IC_{50}$ [µM]
0
5
10
15
20
25
30
40
60
*
*
*
****
****
****
****
*
**
****
****
HT-29
HCT116
Ls-180
SW480
LoVo
SW620
DLD1
Ls1034
FHC
HLMEC
Cell lines

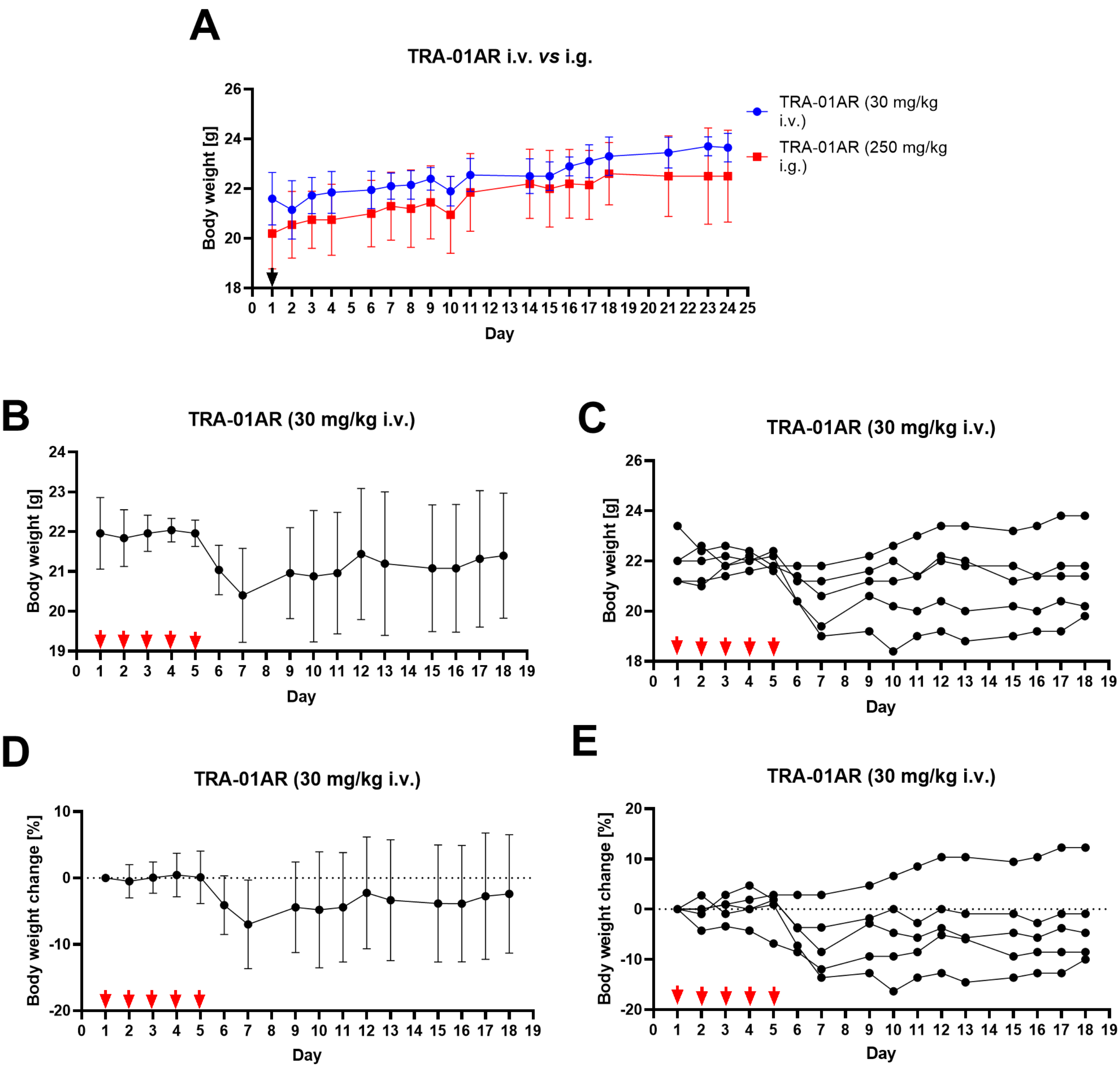
A
TRA-01AR i.v. vs i.g.
Body weight [g]
Day
TRA-01AR (30 mg/kg i.v.)
TRA-01AR (250 mg/kg i.g.)
B
TRA-01AR (30 mg/kg i.v.)
Body weight [g]
Day
C
TRA-01AR (30 mg/kg i.v.)
Body weight [g]
Day
D
TRA-01AR (30 mg/kg i.v.)
Body weight change [%]
Day
E
TRA-01AR (30 mg/kg i.v.)
Body weight change [%]
Day

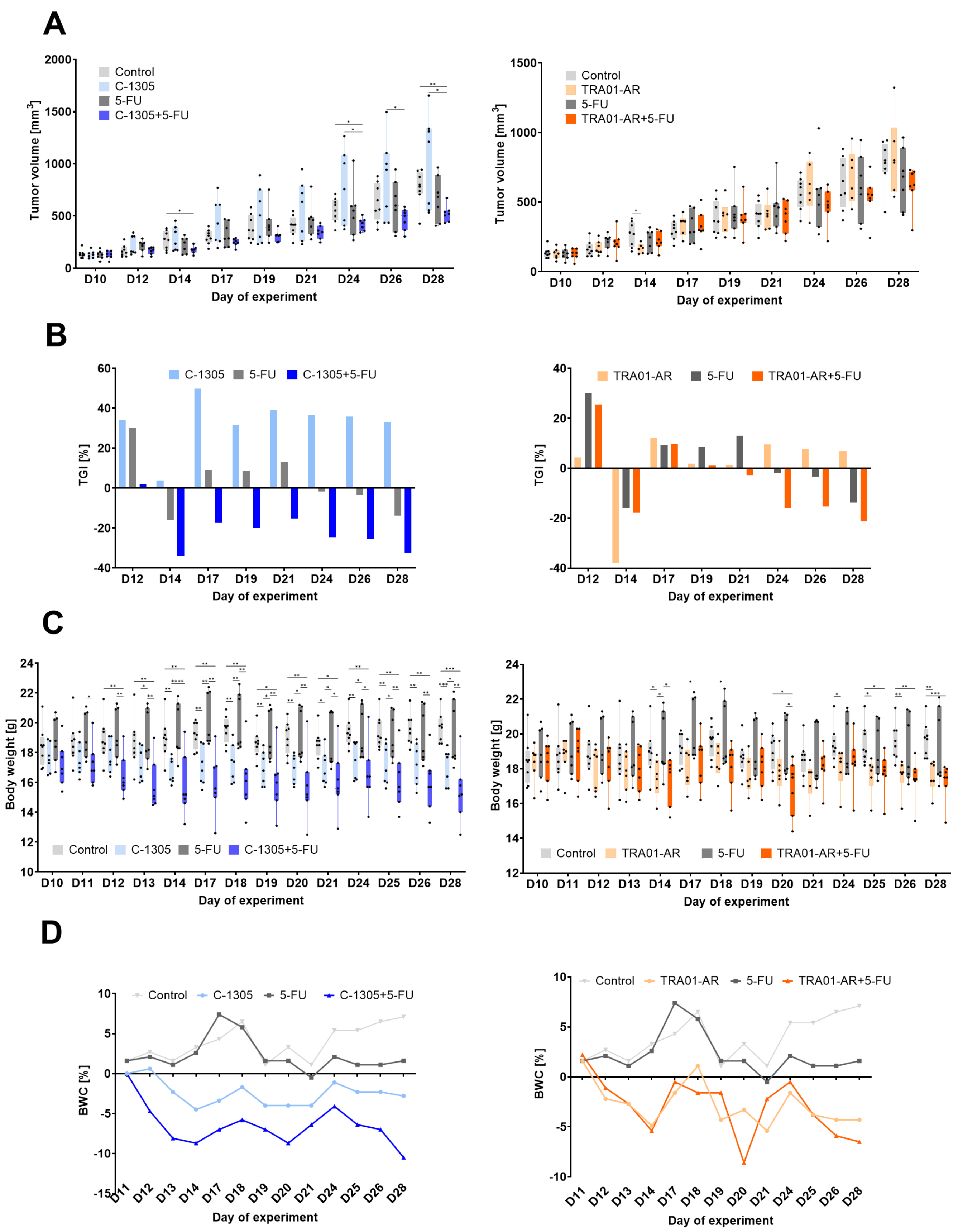
A
Tumor volume [mm3]
Control
C-1305
5-FU
C-1305+5-FU
Day of experiment
TRA01-AR
TRA01-AR+5-FU
B
TGI [%]
C
Body weight [g]
D
BWC [%]
D10 D11 D12 D13 D14 D17 D18 D19 D20 D21 D24 D25 D26 D28

**Supplementary Figure 5**

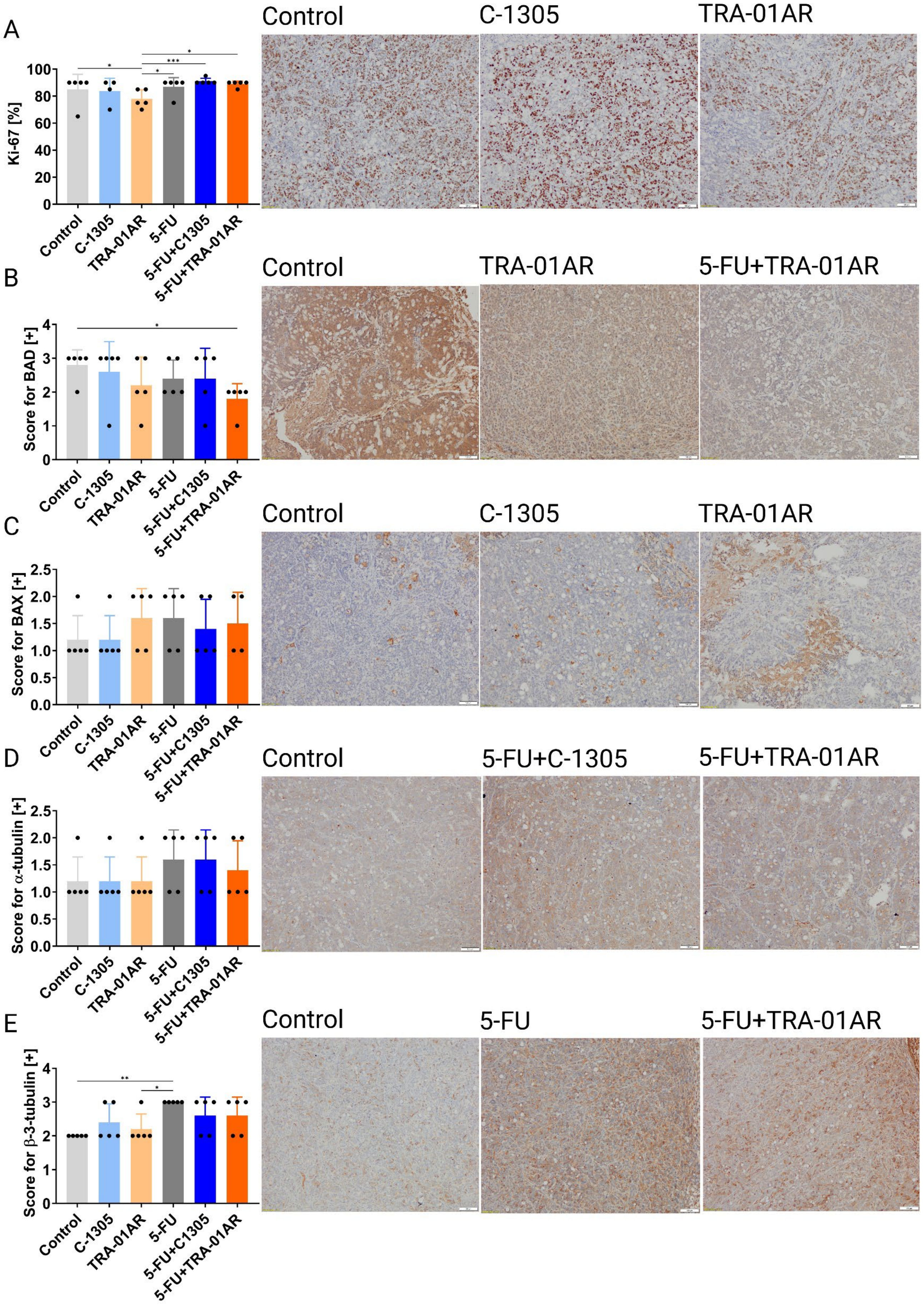

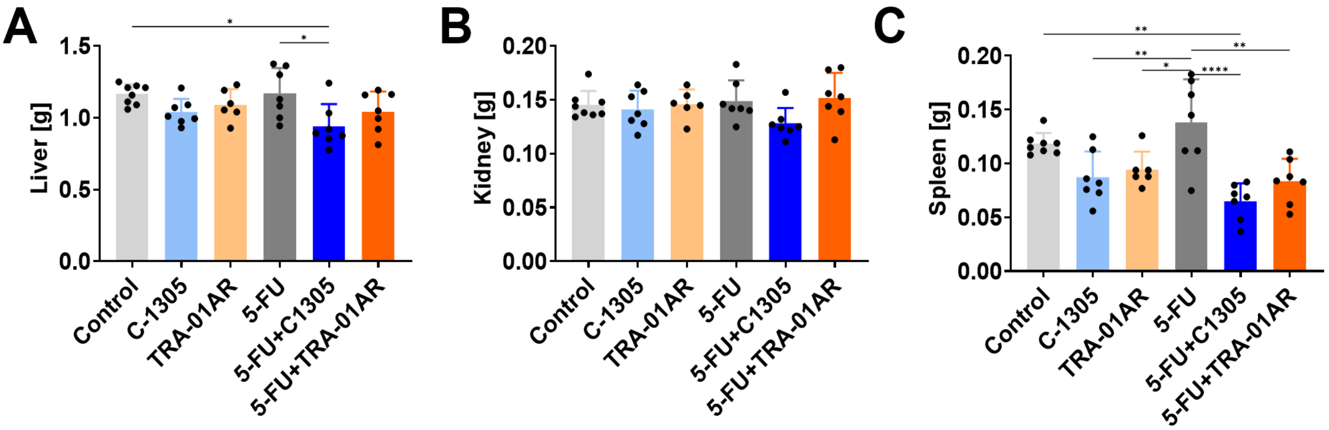
A
Liver [g]
1.5
1.0
0.5
0.0
Control
C-1305
TRA-01AR
5-FU
5-FU+C1305
5-FU+TRA-01AR
B
Kidney [g]
0.20
0.15
0.10
0.05
0.00
C
Spleen [g]

### Supplementary Figure 7

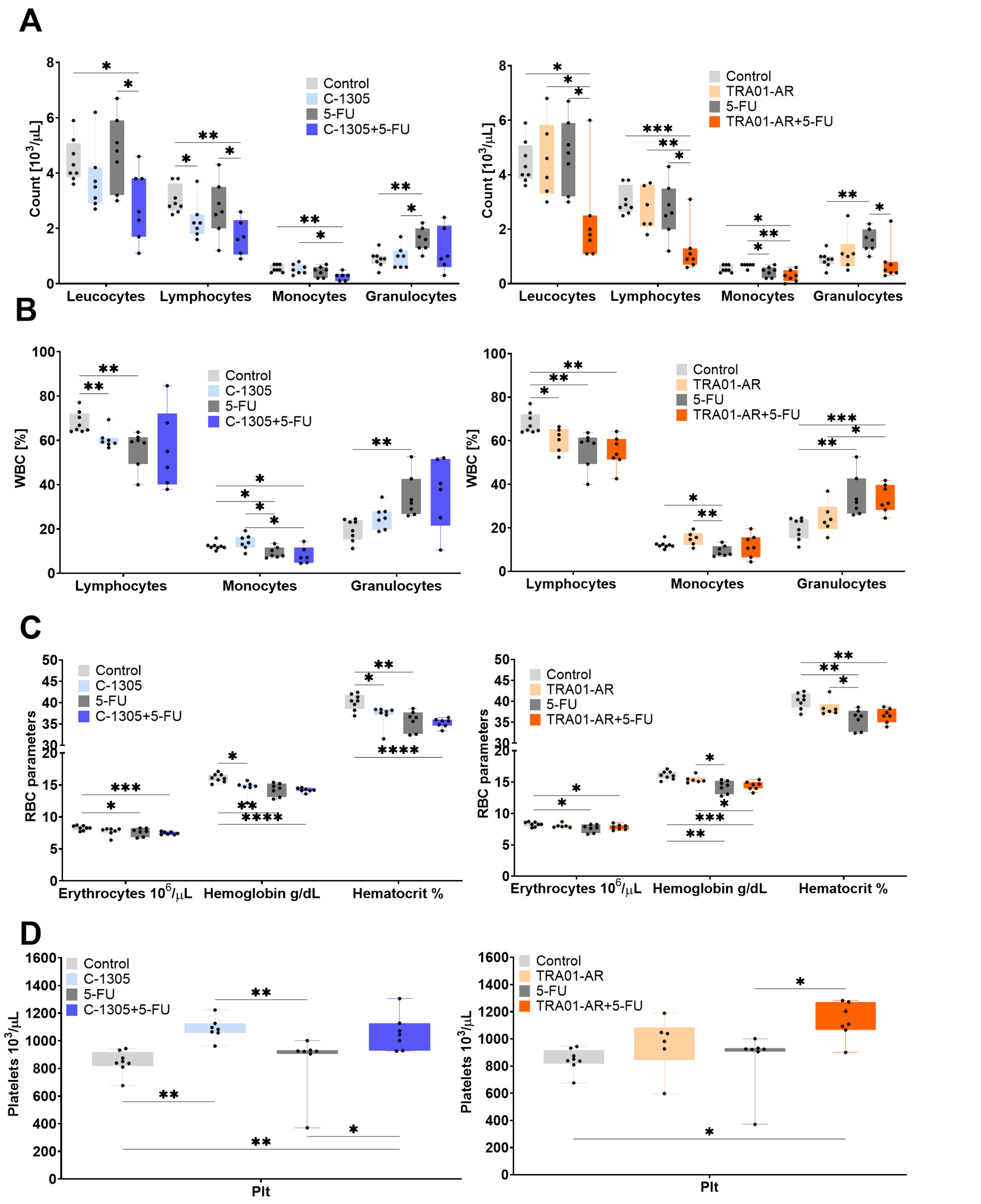

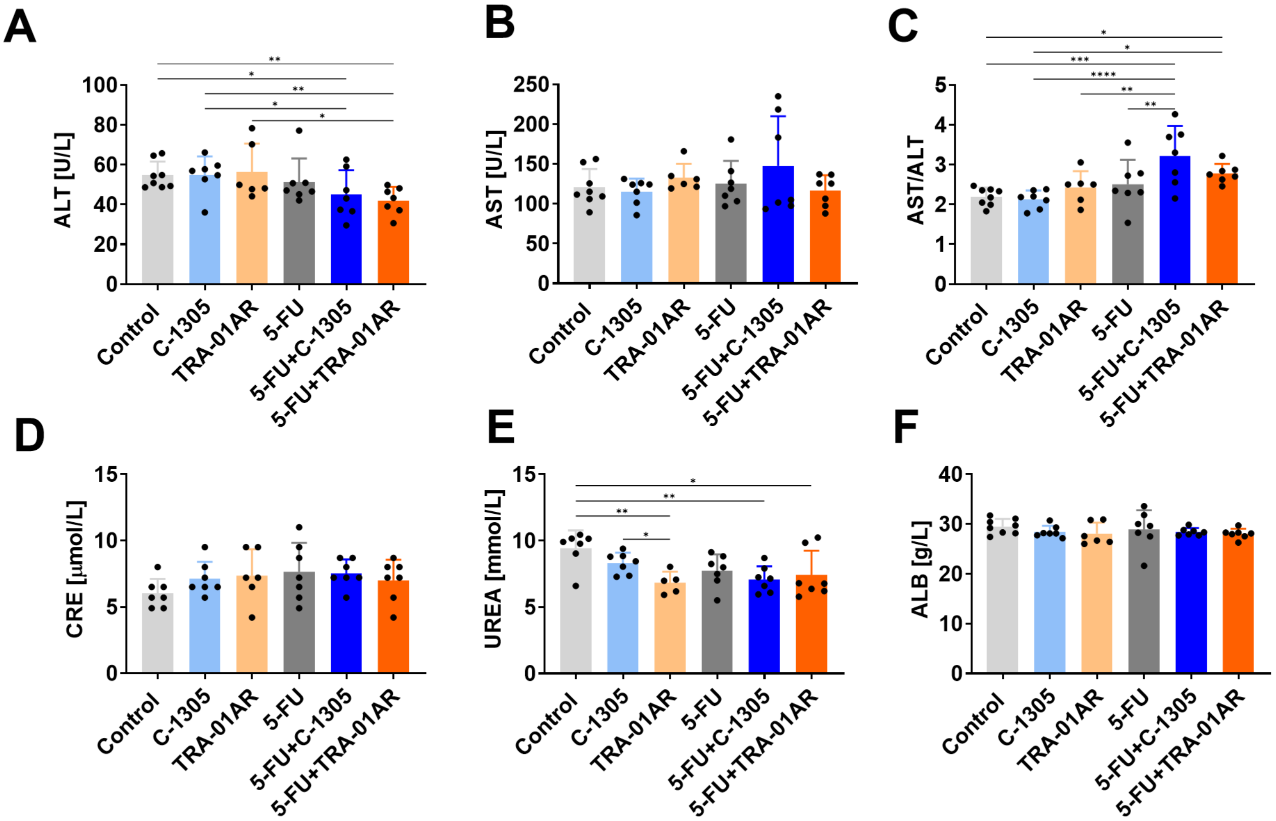
A
ALT [U/L]
Control
C-1305
TRA-01AR
5-FU
5-FU+C-1305
5-FU+TRA-01AR
B
AST [U/L]
C
AST/ALT
D
CRE [μmol/L]
E
UREA [mmol/L]
F
ALB [g/L]
