## Supplementary material for "Preclinical Evaluation of a Novel BOC-Modified Triazoloacridinone (TRA-01AR) in Combination with 5-Fluorouracil in Colorectal Cancer": Table S2

**Supplementary Table 2**: Summary of statistical analyses

| **Figure/Panel** | **Statistical test** | **Effect** | **Statistic: ANOVA: F (DFn, DFd) or Kruskall-Wallis: H** | **p-value** |
| --- | --- | --- | --- | --- |
| Fig.3A 6h-OXPHOS | One-way ANOVA | Treatment | F (3, 15) = 5.929 | =0.0071 |
| Fig.3A 24h-OXPHOS | One-way ANOVA | Treatment | F (3, 19) = 11.36 | =0.0002 |
| Fig.3A 48h-OXPHOS | One-way ANOVA | Treatment | F (3, 19) = 22.72 | <0.0001 |
| Fig.3A 6h glycolysis | One-way ANOVA | Treatment | F (3, 13) = 12.26 | =0.0004 |
| Fig.3A 24h glycolysis | One-way ANOVA | Treatment | F (3, 19) = 11.36 | =0.0002 |
| Fig.3A 48h glycolysis | One-way ANOVA | Treatment | F (3, 19) = 22.72 | <0.0001 |
| Fig 3B 6h-OXPHOS | One-way ANOVA | Treatment | F (3, 12) = 5.762 | =0.0112 |
| Fig 3B 24h-OXPHOS | One-way ANOVA | Treatment | F (3, 19) = 369.0 | <0.0001 |
| Fig 3B 48h-OXPHOS | One-way ANOVA | Treatment | F (3, 19) = 56.78 | <0.0001 |
| Fig 3B 6h-glycolysis | One-way ANOVA | Treatment | F (3, 12) = 325.4 | <0.0001 |
| Fig 3B 24h- glycolysis | One-way ANOVA | Treatment | F (3, 19) = 363.3 | <0.0001 |
| Fig 3B 48h- glycolysis | One-way ANOVA | Treatment | F (3, 19) = 56.78 | <0.0001 |
| Fig. 4A 6h-OXPHOS | One-way ANOVA | Treatment | F (3, 16) = 1.245 | =0.3264 |
| Fig. 4A 24h-OXPHOS | One-way ANOVA | Treatment | F (3, 19) = 6.127 | =0.0043 |
| Fig. 4A 48h-OXPHOS | One-way ANOVA | Treatment | F (3, 19) = 4.176 | =0.0198 |
| Fig 4A 6h-glycolysis | One-way ANOVA | Treatment | F (3, 16) = 0.5297 | =0.6683 |
| Fig 4A 24h-glycolysis | One-way ANOVA | Treatment | F (3, 19) = 6.127 | =0.0043 |
| Fig 4A 48h-glycolysis | One-way ANOVA | Treatment | F (3, 19) = 4.176 | =0.0198 |
| Fig. 4B 6h-OXPHOS | One-way ANOVA | Treatment | F (3, 17) = 0.1728 | =0.9133 |
| Fig. 4B 24h-OXPHOS | One-way ANOVA | Treatment | F (3, 19) = 30.45 | <0.0001 |
| Fig. 4B 48h-OXPHOS | One-way ANOVA | Treatment | F (3, 19) = 24.90 | <0.0001 |
| Fig. 4B 6h-glycolysis | One-way ANOVA | Treatment | F (3, 17) = 0.3257 | =0.8068 |
| Fig. 4B 24h-glycolysis | One-way ANOVA | Treatment | F (3, 19) = 30.45 | <0.0001 |
| Fig. 4B 48h-glycolysis | One-way ANOVA | Treatment | F (3, 19) = 26.52 | <0.0001 |
| Fig. 6A | One-way ANOVA | Treatment | F (5, 36) = 3.798 | =0.0073 |
| Fig. 6B WiPi | One-way ANOVA | Treatment | F (5, 29) = 3.084 | =0.0237 |
| Fig. 6B WiR | One-way ANOVA | Treatment | F (5, 34) = 7.856 | <0.0001 |
| Fig. 6B PE | One-way ANOVA | Treatment | F (5, 34) = 2.979 | =0.0246 |
| Fig. 6B WiAUC | One-way ANOVA | Treatment | F (5, 27) = 0.8788 | =0.5085 |
| Fig. 7 A area of necrosis | Kruskal-Wallis | --- | H=23.38 | =0.0003 |
| Fig. 7 A CD31 | One-way ANOVA | Treatment | F (5, 23) = 2.090 | =0.1033 |
| Fig. 7 A TUNEL | Kruskal-Wallis | --- | H=15.93 | =0.0070 |
| Fig. 7 A TERT | Kruskal-Wallis | --- | H=9.512 | =0.0903 |
| Fig. 8 A IL-6 | One-way ANOVA | Treatment | F (5, 16) = 4.266 | =0.0117 |
| Fig. 8 A CCL2 | One-way ANOVA | Treatment | F (5, 18) = 7.812 | =0.0005 |
| Fig. 8 B FGF1 | One-way ANOVA | Treatment | F (5, 29) = 6.860 | =0.0002 |
| Fig. 8 B FGF2 | One-way ANOVA | Treatment | F (5, 27) = 4.720 | =0.0032 |
| Fig. 8 B VEGF | One-way ANOVA | Treatment | F (5, 27) = 6.277 | =0.0006 |
| Fig. 8 B PDGF | One-way ANOVA | Treatment | F (5, 29) = 2.754 | =0.0373 |
| Fig. 8 B CCL2 | One-way ANOVA | Treatment | F (5, 30) = 1.364 | =0.2656 |
| Fig. 8 B IL-6 | One-way ANOVA | Treatment | F (5, 27) = 1.617 | =0.1894 |
| Fig. 8 B TGF-β | One-way ANOVA | Treatment | F (5, 27) = 12.41 | <0,0001 |
| Fig. 8 C ERK1/2 | One-way ANOVA | Treatment | F (5, 18) = 1.828 | =0.1581 |
| Fig. 8 C p-ERK1/2 | One-way ANOVA | Treatment | F (5, 18) = 1.778 | =0.1683 |
| Fig. 8 C pERK/ERK | One-way ANOVA | Treatment | F (5, 18) = 1.737 | =0.1772 |
| Fig. 8 D COX-2 | Kruskal-Wallis | --- | H=7.328 | =0.1974 |
| Fig. 8 D eNOS | One-way ANOVA | Treatment | F (5, 18) = 2.863 | =0.0450 |
| Fig. 8 D Ang-4 | Kruskal-Wallis | --- | H=2.080 | =0.8380 |
| Fig. 8 D p53 | Kruskal-Wallis | --- | H=2.692 | =0.7473 |
| Suppl. Fig.S4 A left panel | Two-way ANOVA | Interaction | F (4.563, 38.03) = 3.445 | =0.0136 |
| Suppl. Fig.S4 A left panel |  | Time | F (1.521, 38.03) = 124.6 | <0.0001 |
| Suppl. Fig.S4 A left panel |  | Treatment | F (3, 25) = 2.899 | =0.0549 |
| Suppl. Fig.S4 A right panel | Two-way ANOVA | Interaction | F (6.291, 50.59) = 1.800 | =0.1147 |
| Suppl. Fig.S4 A right panel |  | Time | F (2.097, 50.59) = 148.9 | <0.0001 |
| Suppl. Fig.S4 A right panel |  | Treatment | F (3, 25) = 0.2826 | =0.8375 |
| Suppl. Fig.S4 C left panel | Two-way ANOVA | Interaction | F (11.36, 94.70) = 6.185 | <0.0001 |
| Suppl. Fig.S4 C left panel |  | Time | F (3.788, 94.70) = 7.054 | <0.0001 |
| Suppl. Fig.S4 C left panel |  | Treatment | F (3, 25) = 7.409 | =0.0010 |
| Suppl. Fig.S4 C right panel | Two-way ANOVA | Interaction | F (13.71, 110.7) = 4.333 | <0.0001 |
| Suppl. Fig.S4 C right panel |  | Time | F (4.570, 110.7) = 7.726 | <0.0001 |
| Suppl. Fig.S4 C right panel |  | Treatment | F (3, 25) = 2.151 | 0.1191 |
| Suppl. Fig. S5 Ki-67 | Kruskal-Wallis | --- | H=12.49 | =0.0286 |
| Suppl. Fig. S5 BAD | Kruskal-Wallis | --- | H= 6.930 | =0.2259 |
| Suppl. Fig. S5 BAX | Kruskal-Wallis | --- | H= 3.322 | =0.6505 |
| Suppl. Fig. S5 α-tubulin | Kruskal-Wallis | --- | H= 4.024 | =0.5460 |
| Suppl. Fig. S5 β3-tubulin | Kruskal-Wallis | --- | H= 11.91 | =0.0360 |
| Suppl. Fig. S6 A | One-way ANOVA | Treatment | F (5, 36) = 3.277 | =0.0154 |
| Suppl. Fig. S6 B | Kruskal-Wallis | --- | H= 7.965 | =0.1582 |
| Suppl. Fig. S6 C | One-way ANOVA | Treatment | F (5, 36) = 9.144 | <0.0001 |
| Suppl. Fig S7 A left panel | Two-way ANOVA | Interaction | F (5.183, 41.46) = 4.265 | =0.0029 |
|  |  | Cell type | F (1.728, 41.46) = 247.6 | <0.0001 |
|  |  | Treatment | F (3, 25) = 3.603 | =0.0273 |
| Suppl. Fig S7 A right panel | Two-way ANOVA | Interaction | F (4.128, 33.03) = 5.957 | =0.0009 |
|  |  | Cell type | F (1.376, 33.03) = 211.7 | <0.0001 |
|  |  | Treatment | F (3, 24) = 5.157 | =0.0068 |
| Suppl. Fig S7 B left panel | Two-way ANOVA | Interaction | F (3.399, 27.19) = 3.760 | =0.0187 |
|  |  | Cell type | F (1.133, 27.19) = 175.7 | <0.0001 |
|  |  | Treatment | F (3, 24) = 14.67 | <0.0001 |
| Suppl. Fig S7 B right panel | Two-way ANOVA | Interaction | F (4.042, 32.34) = 6.419 | =0.0006 |
|  |  | Cell type | F (1.347, 32.34) = 305.9 | <0.0001 |
|  |  | Treatment | F (3, 24) = 98.79 | <0.0001 |
| Suppl. Fig S7 C left panel | Two-way ANOVA | Interaction | F (3.163, 26.36) = 10.00 | =0.0001 |
|  |  | Parameter type | F (1.054, 26.36) = 9608 | <0.0001 |
|  |  | Treatment | F (3, 25) = 8.359 | =0.0005 |
| Suppl. Fig S7 C right panel | Two-way ANOVA | Interaction | F (3.116, 24.93) = 7.806 | =0.0007 |
|  |  | Parameter type | F (1.039, 24.93) = 9426 | <0.0001 |
|  |  | Treatment | F (3, 24) = 7.232 | =0.0013 |
| Suppl. Fig S7 D left panel | One-way ANOVA | Treatment | F (3, 25) = 6.450 | =0.0022 |
| Suppl. Fig S7 D right panel | One-way ANOVA | Treatment | F (3, 24) = 4.841 | =0.0090 |
| Suppl. Fig. S8 A | Kruskal-Wallis | --- | H=12.05 | =0.0341 |
| Suppl. Fig. S8 B | One-way ANOVA | Treatment | F (5, 36) = 1.010 | =0.4263 |
| Suppl. Fig. S8 C | One-way ANOVA | Treatment | F (5, 36) = 5.632 | =0.0006 |
| Suppl. Fig. S8 D | One-way ANOVA | Treatment | F (5, 35) = 0.9585 | =0.4563 |
| Suppl. Fig. S8 E | Kruskal-Wallis | --- | H=13.85 | =0.0166 |
| Suppl. Fig. S8 F | One-way ANOVA | Treatment | F (5, 36) = 0.5671 | P=0.7246 |
